# Pre-processing of paleogenomes: Mitigating reference bias and postmortem damage in ancient genome data

**DOI:** 10.1101/2023.11.11.566695

**Authors:** Dilek Koptekin, Etka Yapar, Kıvılcım Başak Vural, Ekin Sağlıcan, N. Ezgi Altınışık, Anna-Sapfo Malaspinas, Can Alkan, Mehmet Somel

## Abstract

Ancient DNA analysis is subject to various technical challenges, including bias towards the reference allele (“reference bias”), postmortem damage (PMD) that confounds real variants, and limited coverage. Here, we conduct a systematic comparison of alternative approaches against reference bias and against PMD. To reduce reference bias, we either (a) mask variable sites before alignment or (b) align the data to a graph genome representing all variable sites. Compared to alignment to the linear reference genome, both masking and graph alignment effectively remove allelic bias when using simulated or real ancient human genome data, but only if sequencing data is available in FASTQ or unfiltered BAM format. Reference bias remains indelible in quality-filtered BAM files and in 1240K-capture data. We next study three approaches to overcome postmortem damage: (a) trimming, (b) rescaling base qualities, and (c) a new algorithm we present here, ***bamRefine***, which masks only PMD-vulnerable polymorphic sites. We find that bamRefine is optimal in increasing the number of genotyped loci up to 20% compared to trimming and in improving accuracy compared to rescaling. We propose graph alignment coupled with bamRefine to minimise data loss and bias. We also urge the paleogenomics community to publish FASTQ files.

## INTRODUCTION

Ancient DNA (aDNA) has become today a major information source for studies of evolution or the human past. However, paleogenomic data has its specific challenges, being characterised by short fragment lengths, post-mortem damage (PMD) in the form of transitions at the ends of DNA molecules, and a low abundance of endogenous DNA resulting in low coverage genomes. Standard aDNA data processing pipelines typically involve (i) alignment of reads to a linear reference genome, (ii) quality filtering of reads, (iii) modifications to the read data to avoid PMD confounding with true genetic variation, such as trimming or rescaling, (iv) genotyping at known polymorphic loci (as low coverage generally precludes *de novo* genotyping), (v) pseudohaploidization, i.e., randomly choosing one allele per variant site (a strategy to overcome biases related to heterogeneous coverage among studied genomes). These procedures are susceptible to various biases and shortcomings that can eventually lead to inaccurate interpretations of genetic relationships, population history, or evolutionary processes. We will tackle two of such issues in this study: reference bias, and biased and/or low-efficiency genotyping in the face of PMD.

Although biases against divergent ancient DNA reads had been noted earlier (1), the reference bias phenomenon in ancient genomes was first coined and explained by Günther and Nettelblad (2019). These authors described how read alignment to a linear reference genome with low-coverage and short read-based sequencing data can lead to a higher frequency of reference allele calls over alternative allele calls at heterozygous sites when a 1:1 ratio would be expected. Reference bias arises due to the read alignment quality score calculation: reads with mismatches receive lower scores than perfectly matched reads.

Hence non-reference allele-carrying reads tend to be either unmapped or assigned lower alignment quality scores than the reference allele-carrying reads, and thus removed by filtering reads for by a minimal alignment quality score. Consequently, reference allele-carrying reads are overrepresented in the aligned and filtered data. Reference biases have been observed to impact population genetic and phylogenetic analyses of present-day taxa when evolutionarily distant linear reference genomes are used for alignment (2,3).

Meanwhile, ancient DNA sequencing data is particularly prone to such bias, because when reads are short and/or have higher residual PMD, mismatches caused by alternative alleles can have a disproportionate impact on quality scores. The overrepresentation of reference allele-carrying reads may render ancient genome profiles more similar to the reference genome than they actually are. This effect can then lead to biased results in downstream inferences on phylogenetics, demographic history or kinship.

Previous studies have suggested several methods to reduce reference bias in ancient DNA studies: (a) Statistically accounting for possible reference bias during variant calling (4), which can be effective but only on high-coverage genomes; (b) aligning reads to a modified version of the linear reference genome, e.g. by representing both alleles or a third allele at variable sites (5–7); (c) modifying ancient reads at variable sites by converting them to ‘N’ (5); (d) using a graph reference genome that represents the variants in large genomic variation datasets such as the 1000 Genomes Project (8).

A second challenge in paleogenome data pre-processing involves ensuring that PMD on molecules does not impact inferred genotypes. One correction strategy is experimentally removing PMD after DNA extraction using uracil-DNA glycosylase (UDG) treatment (9). The majority of researchers who use UDG employ the half-UDG protocol, which still leaves a slight excess of transitions at molecule ends (10). PMD may also be accounted for using post-alignment *in silico* approaches. One solution involves limiting analyses transversions alone, which are much less (only indirectly) affected by PMD (1). However, this approach leads to the loss of approximately 60% of polymorphism data in humans and other mammals, as transition polymorphisms are about twice more numerous than transversions. An alternative, and currently the most prevalent method is trimming, or masking the end of the reads in a BAM file. This involves changing bases at read ends of a specific length to ‘N’ and their quality to ‘!’ (corresponding to zero in Phred+33 encoding), e.g. using the tool *trimBAM* (11). Most researchers remove two to three bases at read termini of half-UDG-treated libraries, or 8-10 bases of non-UDG-treated libraries (12). This trimming process also leads to data loss, especially for the latter type of libraries. For instance, in a non-UDG-treated and paired-end library, 10 bp are masked from both ends (2x10=20bp in total) per standard 60 bp aDNA read, which means c.30% data loss. Other methods, such as *mapDamage* (13) and *ATLAS* (14), have attempted to reduce the effect of PMD by rescaling the base quality of possible PMD-driven misincorporations, but such approaches are rarely used as they could alter genotype frequencies, which has not yet been systematically investigated. Yet an alternative approach could be masking only PMD-sensitive regions on read ends, thus retaining more genetic information and enabling more comprehensive analysis of low-coverage ancient genomes.

In this work, we study solutions to reduce the effect of reference bias and PMD. We first investigate the degree of reference bias using linear mapping, mapping to a masked genome, and using a graph genome on simulated as well as real paleogenomic data of various types. We then study genotyping efficiency under PMD using standard trimming, using *mapDamage*, and masking read ends that overlap with genomic positions that are sensitive to PMD-related false positive variant calls using a new algorithm, ***bamRefine***. Our results show that using alternative reference genomes (either graph or masked) together with *bamRefine* results in more accurate genotypes and reduces data loss.

## MATERIALS AND METHODS

### Simulating ancient genomes

We used chromosome 1 of the human reference genome (version hs37d5) as a template to generate the simulated ancient genome data. We chose bi-allelic SNPs on chromosome 1 of the individual 06A010111 of the Turkish Genome Project dataset (15), which consisted of 182,515 homozygous reference, 53,391 homozygous alternative, 77,841 heterozygous positions (313,747 positions in total) (see Table S1). We then inserted these into the chromosome 1 template with “*vcftools/vcf-consensus (v.0.1.6)*” (16).

We generated ancient DNA data using “*gargammel*” (17), using the template chromosome 1 data with polymorphism inserted. Five “*gargammel*” simulations were performed for five target coverages: 0.05X, 0.1X, 1X, 5X and 10X. We used a normal distribution with a mean of 65 bp for the read size distribution. The parameter “*-damage 0.024,0.36,0.0097,0.55”* was used to introduce PMD to simulated ancient genomes (see Figure S1). We did not include bacterial or modern contamination in the data by using the “*-comp 0,0,1*” parameter.

### Real ancient genomes

We selected 17 published ancient genomes, either shotgun-sequenced, whole-genome captured or 1240K SNP-captured, all from human skeletal material originating from different geographic regions (12,18–31). The coverage of samples ranges from low to medium coverage to high coverage. The dataset includes both damage-repaired and non-damage-repaired samples (see Table S2).

The raw FASTQ files of 7 out of 17 samples were available. Others were downloaded as BAM files and converted to FASTQ files using “*Picard SamToFastq (version 2.23.8)*” (http://broadinstitute.github.io/picard/). A number of FASTQ files were not publicly available (Table S2) and were provided by the research teams upon request.

### Alignment strategies

We removed the residual adapter sequences in raw FASTQ files for each sample using the software ‘‘*Adapter Removal (version 2.3.1)”* (32) using ‘‘*–qualitybase 33 –gzip –trimns*’’ parameters. The reads in paired-end libraries were merged after removing residual adapter sequences, requiring at least 11 bp overlap between the pairs using the additional parameter ‘‘*–collapse –minalignmentlength 11*’’.

We aligned FASTQ files to three different reference genomes:

1. <u>Linear Reference Genome (version hs37d5):</u> We used the program ‘‘*BWA aln/samse (version 0.7.15)*” (33) with parameters ‘‘*-n 0.01, -o 2’*’ and disabled the seed with ‘*‘-l 16500*’.
2. <u>Masked Linear Reference Genome (masked version of hs37d5):</u> We masked the positions we wanted to genotype on the linear reference genome using “*bedtools maskfasta (v. 2.29.1)*” (34) by converting the nucleotides to ‘N’. After masking, we aligned samples using ‘‘*BWA aln/samse (version 0.7.15)”* (33) with the same parameters above.
3. <u>Graph Reference Genome:</u> We obtained a published graph genome version from Seven Bridges Inc. (*SBG.Graph.B37.V6.rc6.vcf.gz*), which included variants from 1000 Genomes

(1000G) Phase 3 (with alternate allele frequency greater than 0.01) (35), the Simons Genome Diversity Panel (alternate allele occurrence of 10 or greater) (36), and other INDEL variant datasets to construct a graph genome. We used the “*GRAF tool (version 0.12.5)*” (37) to align the reads to this graph genome annotation together with the baseline reference genome GRCh37. See (37) and https://www.sevenbridges.com/graph-genome-academic-release/ for more details.

After alignment, we removed PCR duplicates using “*FilterUniqueSAMCons.py”* (38) and removed reads <35 bp, with >10% mismatches to the reference genome, or with <30 mapping quality (MAPQ) from all BAM files.

We added read group information to final BAM files by using “*picard AddOrReplaceReadGroups (version 2.23.8)*” (http://broadinstitute.github.io/picard/).

### bamRefine

Here, we present a new variant-aware PMD-correction algorithm called *bamRefine. bamRefine* is an efficient tool to prevent possible PMD-affected bases at the read ends from being included in the variant calling process based on a given variant list. It has a simple algorithm with two main steps: First, the variant list to be used in downstream analyses is parsed and classified into 5’ and 3’ “suspect” lists, which in this context corresponds to variants with “C” and “G” alleles, respectively. “Suspects” here refer to the genomic positions that carry the risk of “C->T” or “G->A” false-positive variant calls if PMD-affected bases at read ends were to be used without any form of masking in downstream steps of a pipeline.

Then, the BAM file is processed read by read, masking bases that overlap with 5’ (3’) suspects within a user-specified nucleotide lookup range from the 5’ (3’) end of each read. The lookup range is determined based on the PMD signature in the library. The program allows the 5’ and 3’ end lookup ranges to be asymmetrical to properly process reads from single-stranded library protocols. The masking is confined to the positions that overlap with the 5’/3’ variant tables within the user-specified lookup range from 5’/3’ ends of reads and is implemented regardless of the allele an individual read carries (Figure 1). This results in less data loss when compared to trimming the entire lookup range (e.g. T/G variants are retained at 5’ ends) and ensures non-biased masking (e.g. PMD can shift allele frequencies at C/G variants at both ends and masking these avoids this effect). The job of flagging and masking positions of interest for each chromosome in a BAM file is parallelized by multiprocessing, allowing the program to rapidly refine millions of reads. More detailed information regarding usage and installation instructions can be found at https://github.com/etkayapar/bamRefine. *bamRefine* is also implemented in the *Mapache* ancient DNA pre-processing pipeline (39) (https://github.com/sneuensc/mapache).

**Figure 1:**
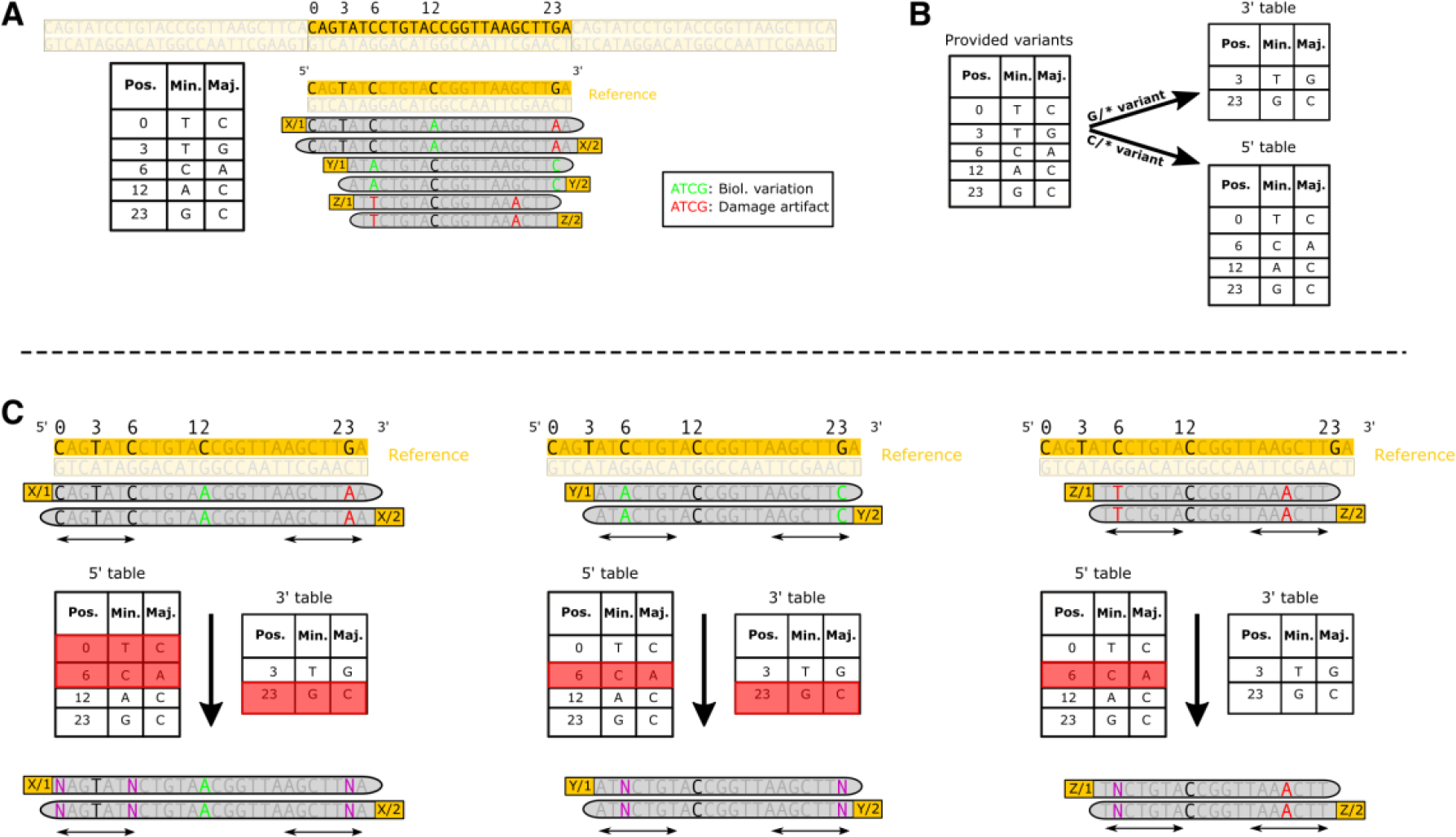
Graphical representation of the bamRefine workflow run with a BAM file with 6 reads, a variant list with 5 variants, and using the “*--pmd-length-threshold 7”* parameter (i.e. a lookup range of 7 bp at each read end). **A)** A cartoon genome browser view of all the reads mapped to the genomic region, and the input variant list shown as a table. **B)** The classification of the provided variant list into 5’ and 3’ suspects. **C)** Masking of the 6 different reads according to the specified options and input variant list. Masking happens regardless of the alleles the reads carry and only depends on a base within the lookup range overlapped with the variant table of the respective read end.

### PMD-correction strategies

We processed the data using three alternative strategies for avoiding the impact of PMD on genotypes.

i. <u>TRIMMING:</u> We applied trimming (clipping) to the sequencing data using the “*trimBam*” algorithm implemented in “*bamUtil (version 1.0.14)*” (11). We trimmed (a) 10 bases from the ends of each read in non-UDG-treated samples as well as in simulated ancient genomes, and (b) 2 bases from the ends of each read in UDG-treated samples.
ii. <u>RESCALING:</u> We applied rescaling to the sequencing data using the “*mapDamage2”* software (13). We rescaled 10 bases from the ends of each read in simulated data using “*-- rescale --seq-length 10*” parameters. We were unable to execute *mapDamage* analysis on UDG-treated real ancient samples, so we opted not to incorporate a *mapDamage* comparison in our analysis of real ancient data.
iii. <u>REFINING:</u> We applied refining to the sequencing data using “*bamRefine*”. Similar to TRIMMING, we refined (a) 10 bases (using “*--pmd-length-threshold 10*”) from the ends of each read in non-UDG-treated samples as well as the simulated ancient genomes, and (b) 2 bases (using “*--pmd-length-threshold 2*”) from the ends of each read in UDG-treated samples. Regardless of the samples being treated with UDG or not, we used our SNP dataset generated from the Turkish Genome Project for refining the simulated ancient reads (using “*--snps <TGP-SNPS-FILE>* parameter”) and the 1000G sub-Saharan African dataset for the real ancient reads (using “*--snps <AFR-SNPS-FILE>*” parameter).

### Dataset

In previous work we had created a 1000 Genomes sub-Saharan African SNP diversity panel as a high-quality and relatively unbiased SNP dataset to use in demographic inference in Eurasian genomes (12). The dataset includes 4,771,930 (4.7M) bi-allelic autosomal SNPs ascertained in five sub-Saharan African populations in phase 3 of the 1000 Genomes project (35). We used this dataset for genotyping the real ancient genomes included in the analysis.

### Genotyping

We genotyped only targeted SNP positions; for simulated ancient genomes these were the 313,747 positions defined from one individual of the TGP dataset (15), and for real ancient genomes these were the 4.7M positions from 1000 Genomes sub-Saharan African dataset (12). We called both diploid and pseudohaploid genotypes.

<u>Diploid genotypes:</u> These were obtained using “*GATK HaplotypeCaller (version 4.0.11.0)*” (40) by using the “*--min-base-quality-score 30, --minimum-mapping-quality 30, --genotyping-mode GENOTYPE_GIVEN_ALLELES, --output-mode EMIT_ALL_SITES*” parameters as well as the “*--alleles*” parameter to genotype the list of targeted SNP positions.

<u>Pseudo-haploid genotypes:</u> These were obtained by using “*pileupCaller (version 1.4.0)*” (https://github.com/stschiff/sequenceTools) by selecting one allele for each of the targeted SNP positions from the “*samtools mpileup*” (41) output file, which was generated by using the *“-R -B -q30 -Q30”* and the *“-l”* parameters to genotype the list of targeted SNP positions.

### f_4_ statistics

We calculated f_4_-statistics by using “*qpDstat (version: 980)*” algorithm implemented in “*AdmixTools (version 7.0.2)*”(42). We used tests of the form f_4_(Human Reference Genome, Outgroup; Ind1_MappingStrategy1, Ind1_MappingStrategy2) or f_4_(Human Reference Genome, Outgroup; Ind1_MappingStrategy1_PMDCorrectionStrategy1, Ind1_MappingStrategy1_PMDCorrectionStrategy2) using the Chimp Reference Genome (version panTro6) as an outgroup and with the *“f4mode: YES”* option. We used >10,000 overlapping SNPs as cut-off for reporting f_4_-test calculations.

### Visualisation

We produced all graphs in R (43) after reading and manipulating data using “*gsheet”* (44) and “*tidyverse*” (45) packages. All figures were produced by using “*ggplot2*” (46) and its extension packages “*ggpubr*” (47), “*ggh4x*” (48) and “*ggpattern*” (49). The multiple panel figures are combined by using the “*patchwork*” package (50). In some figures, colours were assigned by using “*MetBrewer*” package (51).

## RESULTS

### Simulated genomes: mapping to masked or graph genomes mitigates reference bias

We first simulated ancient human-like sequencing data to gauge reference bias under various alignment strategies. We used the human chromosome 1 (version hs37d5) reference sequence and 77,841 heterozygous sites chosen from a bi-allelic SNP set from the Turkish Genome Project dataset (15) (see Table S1). We created aDNA-like read data with the *gargammel* tool (17) such that reads would carry either allele at heterozygous sites with equal probability. We produced five such datasets with coverages from 0.05X to 10X with PMD damage (Figure S1). We then aligned this data to reference genomes using three different strategies: (i) the “LINEAR” strategy, which is the standard procedure of mapping to a linear reference genome using *bwa aln* with “*-l 16500 -n 0.01 -o 2*”; (ii) the “MASKED” strategy, where, before alignment with *bwa aln*, we masked the linear reference genome sequence at variable positions to be genotyped by converting those bases to ‘N’; and (iii) the “GRAPH” strategy, where we used a graph reference genome representing both reference and alternative alleles at known variable sites and used *GRAF aligner* for mapping (37). We then randomly called pseudohaploid genotypes at the 77,841 heterozygous sites and calculated the alternative allele proportion. We repeated these last steps 100 times.

In the absence of reference bias, we expect ∼50% of pseudohaploid genotypes at heterozygous positions to represent the alternative allele. However, using the “LINEAR” strategy, we observed consistently lower match rates to the alternative allele across all coverages, i.e. reference bias (48.2-50.4%; on average ∼1% lower than expected; binomial test p<0.0001) (Figure 2A, Tables S3-4). Using the “MASKED” or “GRAPH” strategies instead, we observed either slight or no bias towards either allele: the average fraction of alternative alleles was 50.1-50.3% with the former and 49.8-50.1% with the latter (Figure 2A, Tables S3-4). The deviations from 50% using the latter two strategies were also systematically lower than using the “LINEAR” strategy (Mann-Whitney U test p<0.0001; Figure 2A, Figure S2).

**Figure 2:**
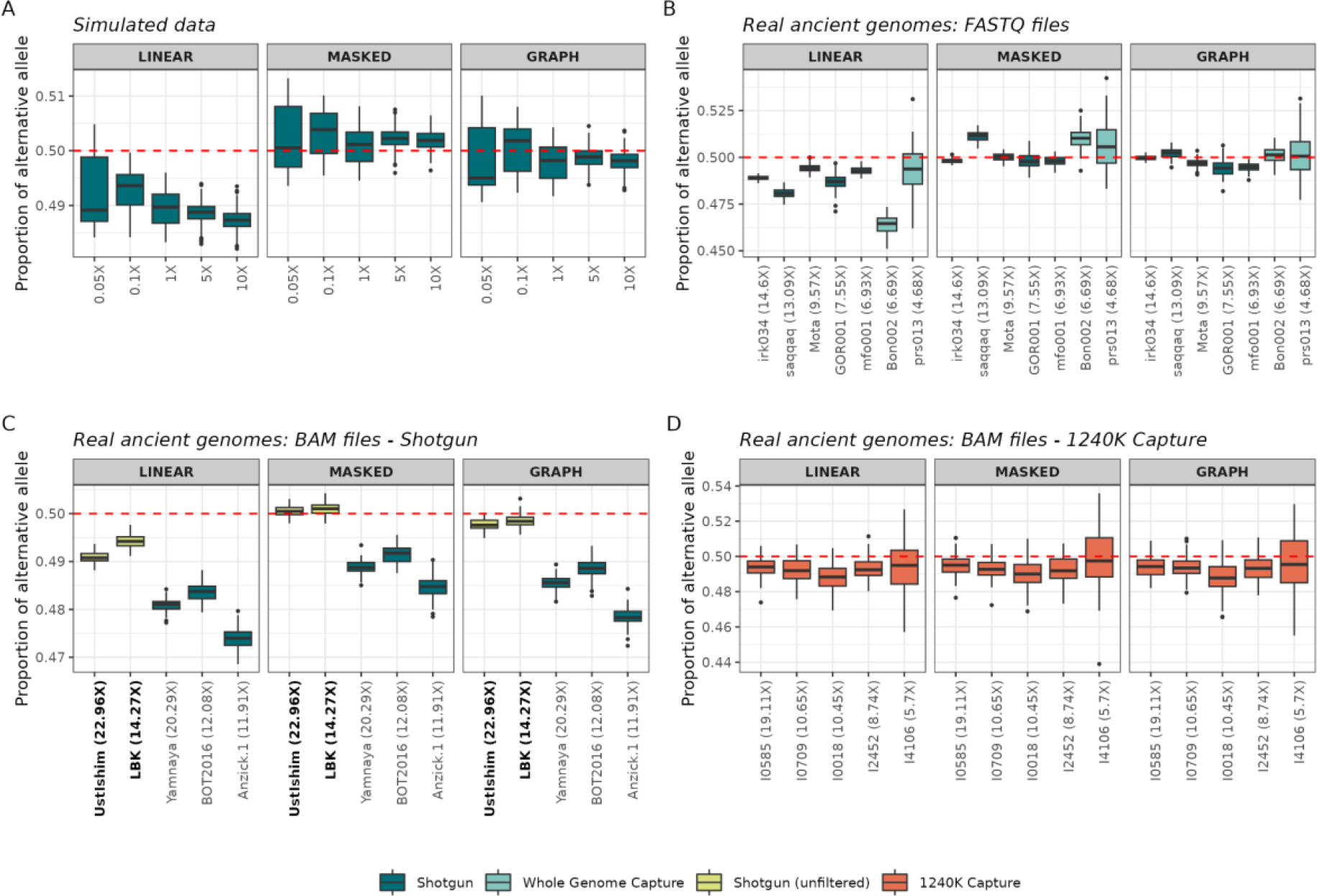
Comparing reference bias under three different alignment strategies (A) using simulated aDNA-like genomes, (B) using real ancient genomes with available raw FASTQ files, and using real (C) shotgun and (D) 1240K capture ancient genomes with already processed BAM files. The plot shows the proportion of alternative alleles after randomly selecting one allele from heterozygote sites 100 times using pileupCaller (panel A: 77,841 sites; panel B: 4,658 - 422,046 sites; panel C: 96,917 - 543,495; panel D: 2,934 - 19,394 sites) (see also Tables S2-6 and Figures S5-6). The BAM files available without strict filtering (i.e., included reads with MAPQ<30) are shown in bold in Panel C (Table S2). In these comparisons we did not apply any PMD-correction.

### Real ancient genomes: reference bias mitigated using FASTQ files but not using BAM files

We next studied reference bias in real paleogenomic data. For this, we started by collecting seven published genomes for which we could obtain raw data as FASTQ files (Table S2). These were derived from diverse geographic regions, produced with or without UDG-treatment, shotgun-sequenced or whole-genome captured, and had variable coverages (Table S2).

We first defined heterozygous sites for each ancient genome as those with 25-75% of reads representing the alternative allele, covered at least by 10X depth and no greater than two times the genome mean coverage (Methods, Table S1). We mapped reads using the three strategies and randomly sampled reads 100 times at these presumed heterozygous sites.

We found salient reference bias using the “LINEAR” strategy, with the fraction of alternative alleles ranging between 46.4-49.4% (binomial test p<0.0001) (Figure 2B, Table S5-6).

Consistent with the simulated data results, the fraction of alternative alleles was ∼50% when using either the “MASKED” (49.8-51.1%) or “GRAPH” strategies (49.4-50.2%) (Mann-Whitney U test p<0.0001; Figure 2B, Figure S3A, Tables S5-6). However, we also noted slight differences between these two approaches: three genomes (Mota, mfo001, GOR001) processed using the “GRAPH” strategy still exhibited a bias against the alternative allele (∼49.5%), whereas two other genomes (Bon002 and Saqqaq) processed using “MASKED” exhibited a weak but significant bias (∼51%) towards the alternative allele (p<0.0001).

Although we lack an explanation for this variability among genomes, we overall conclude that “MASKED” or “GRAPH” approaches both reduce the impact of reference bias on called ancient genotypes (Figure 2A, Figure S3A, Tables S5-6).

The majority of paleogenomes over the last decade have been published as processed BAM files rather than raw FASTQ files, where the former could be subject to irreversible reference bias introduced by mapping parameters as well as alignment quality filtering. To investigate this, we collected 10 additional paleogenomes available as BAM files (Table S2). These included 5 shotgun-generated and 1240K SNPs-enriched genomes. Among the shotgun-generated genomes, the Ust-Ishim and LBK BAM files were published without strict filtering

(i.e., included reads with MAPQ<30), while the rest had been quality filtered (all reads with MAPQ>30) (Figure S4). None of the 1240K SNPs-enriched genomes had been subjected to strict filtering (Figure S4, Table S2).

We again remapped the reads and called pseudo-haploid genotypes using the three strategies. This revealed persistent reference bias for three shotgun-generated BAM files subjected to strict filtering, irrespective of the alignment strategy used (Figure 2C, Figure S4, Table S2). In contrast, both the “MASKED” and “GRAPH” strategies significantly reduced reference bias on Ust-Ishim and LBK, which had not been filtered (Figure 2C, Figure S3B, Table S2). This confirms the expectation that quality filtering of BAM files introduces irreversible reference bias. Meanwhile, all five 1240K SNPs-enriched genomes showed the same level of reference bias irrespective of the alignment strategy used (the alternative allele on average ∼0.7% lower than expected) (Figure 2D, Figure S3B, Figure S4, Table S2). Such bias appears independent of the mapping/filtering process and is likely attributable to 1240K SNP capture favouring one allele over another at targeted SNPs, as reported recently (52,53).

We further observed that the number of reads with MAPQ>30 using the “GRAPH” approach was higher (4 - 21%) than the two other alignment approaches; this was true for all seven FASTQ files (random expectation for “GRAPH” being best in all seven cases: (1/3)^7^ = 0.0004) (Figure S7, Table S7). This happens because reads carrying both alleles would receive higher quality mapping scores using the graph genome alignment than “MASKED” or “LINEAR”. However, we did not observe the same pattern for BAM files, where some of the reads carrying alternative alleles were probably already filtered out.

### Trimming, rescaling and refining as alternative PMD-correction approaches

We also investigated the performance of several approaches for PMD-correction on called genotypes: (A) using “TRIMMING”, i.e. the standard 2 or 10 bp masking of aligned reads with *trimBam* (11), (B) “RESCALING”, which involves rescaling base qualities using *mapDamage2* (13), and (C) “REFINING”, i.e. masking bases at the read ends that overlap with variants sensitive to PMD-related genotyping errors using the new software we present here, ***bamRefine***. Our algorithm masks 5’ end bases only if they overlap with variants that include a “C” allele (to prevent C->T false-positives), or 3’ end bases if they overlap with variants that include a “G” allele (to prevent G->A false-positives). Notably, our approach avoids biased genotyping due to PMD-induced C/G loss at transversion sites (e.g. C’s being underrepresented at a C/A variant site when the variant occurs on 5’ ends of reads due to PMD-induced C->T transitions). Our approach further avoids comprehensive data loss compared with “TRIMMING”, as the latter involves masking extended regions at read ends for non-UDG-treated libraries (Methods).

We used the same simulation scheme using chromosome 1 polymorphisms as above, with the difference that here, along with the 77,841 heterozygous positions described earlier, we also genotyped 182,515 homozygous reference and 53,391 homozygous alternative positions, totalling 313,747 SNPs. We generated aDNA-like read data at 10X coverage using *gargammel* (17), aligned these using either of the three mapping strategies (“LINEAR”, “MASKED”, and “GRAPH”) and applied either of the three PMD-correction approaches mentioned above. We then called diploid genotypes at the 313,747 SNPs using *GATK HaplotypeCaller* (40), and examined the missingness and error rates on these calls, comparing the three PMD-correction approaches.

### Trimming causes data loss and rescaling causes reference bias

Irrespective of the mapping approach, “TRIMMING” exhibited the highest missingness (0.48-0.95%), followed by “REFINING” (0.38-0.86%), and “RESCALING” (0.26-0.52%) (Figure 3A). “TRIMMING” also showed the highest overall error rate among the three methods (2.42-2.48%) (Figure 3A, Figure S5 and Table S4). The bulk of these errors were caused by misassigning heterozygous sites as homozygous reference or homozygous alternative, due to sampling error (i.e. insufficient data to call heterozygous sites) (Figure 3B). “RESCALING” and “REFINING” had slightly lower error rates (2.14-2.31% and 2.07-2.15% respectively), as they use more data than “TRIMMING”. These error rates were higher (∼7%, 6% and 5%, respectively) when repeating the analysis with 5X coverage data (Figure S8); this is expected as with lower coverage sampling error becomes more dramatic.

**Figure 3:**
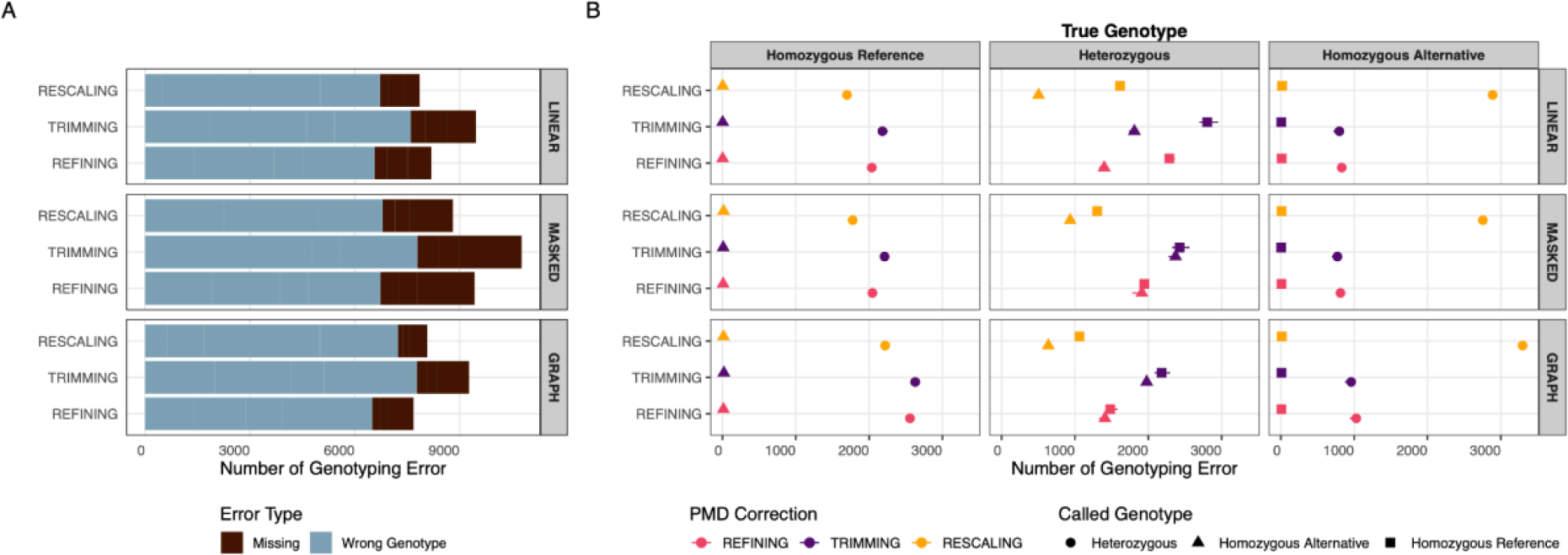
(A) Proportion of genotyping errors and missingness (B) Frequency of type of genotyping errors for each PMD-correction method for 10X coverage simulated genomes, calculated by comparing diploid calls with true genotypes (see also Figures S8-12).

Despite the lowest overall error rate and missingness using “RESCALING”, closer inspection revealed that this approach suffers from significant reference bias. The majority of errors observed with “RESCALING” were caused by favouring the reference allele in genotype calls during PMD-correction (Mann-Whitney U test p<0.0001; Figures S2 and S9), leading to an overestimation of homozygous reference alleles and underestimation of homozygous alternative alleles (Figure 3, Figure S10). In contrast, the “TRIMMING” and “REFINING” approaches label genotypes incorrectly as homozygous reference or alternative at similar rates (Figures S11-12). This pattern could also be observed when calling pseudohaploid genotypes at the 77,841 heterozygous sites after using either of the three PMD-correction methods (Figure 4). “RESCALING” led to a marked underestimation of alternative allele proportions (45.84 - 47.72%), whichever alignment strategy was employed. When we checked transition and transversion sites separately, we observed that this skew in the alternative allele proportions mostly affected transition sites, indicating that the *mapDamage* algorithm used in “RESCALING” overrepresented reference alleles (Figures S13-14).

**Figure 4:**
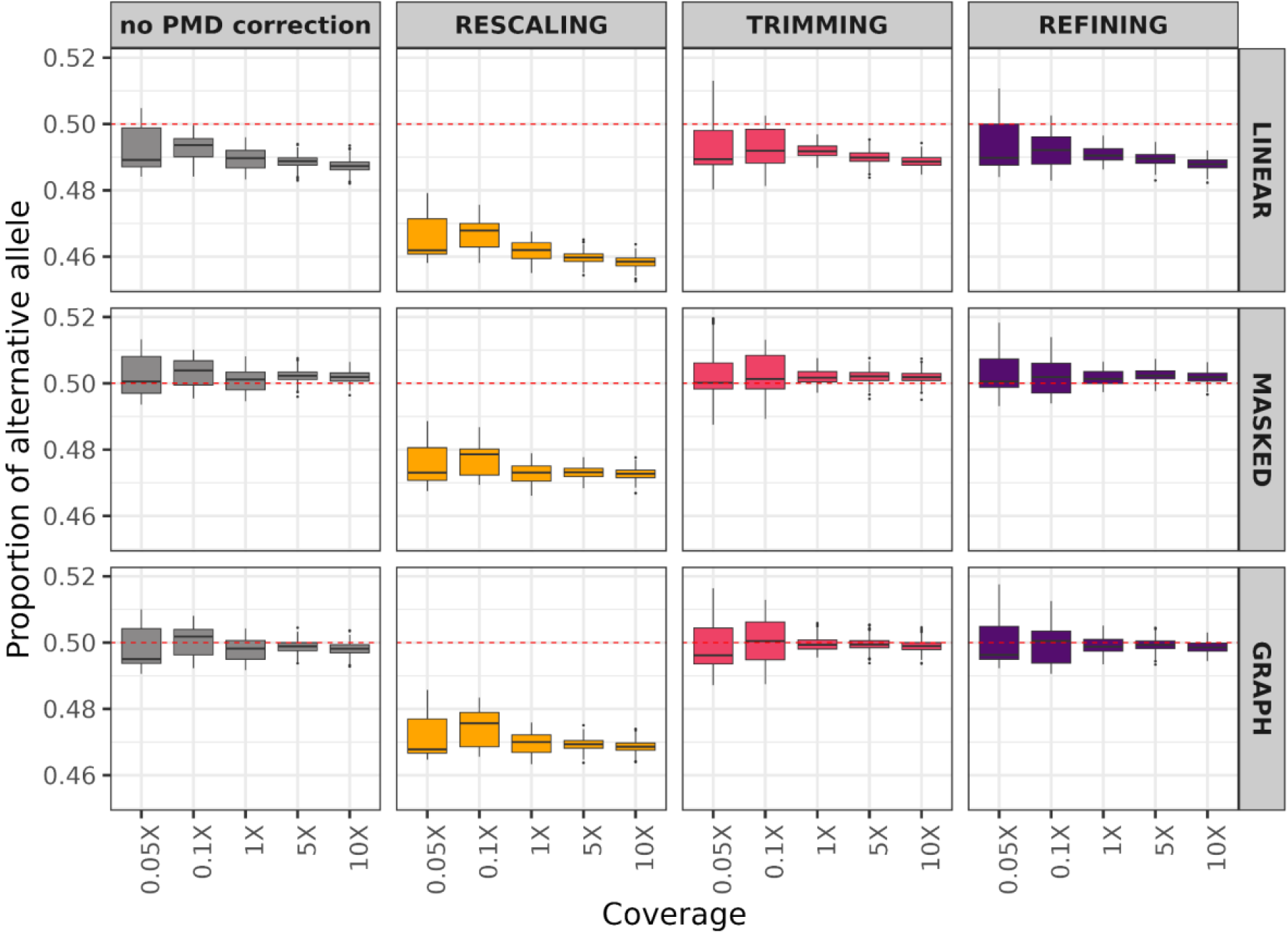
Comparing reference bias in simulated ancient genomes aligned with different reference genomes and PMD-effects reduced with different approaches. The plot shows the proportion of alternative alleles after randomly selecting one allele from heterozygote sites 100 times by *pileupCaller* (see also Figures S13-14).

### Graph or masked mapping followed by bamRefine yields the best F-scores

To investigate genotype accuracies further among the three PMD-correction algorithms, we calculated concordance rate (CR), proportion of false negative, proportion of false positive, non-reference true positive rate (NTPR), as well as recall (or sensitivity) and the F-score (Figure 5), with the alternative allele as our pivot (54) (Figure S15).

**Figure 5:**
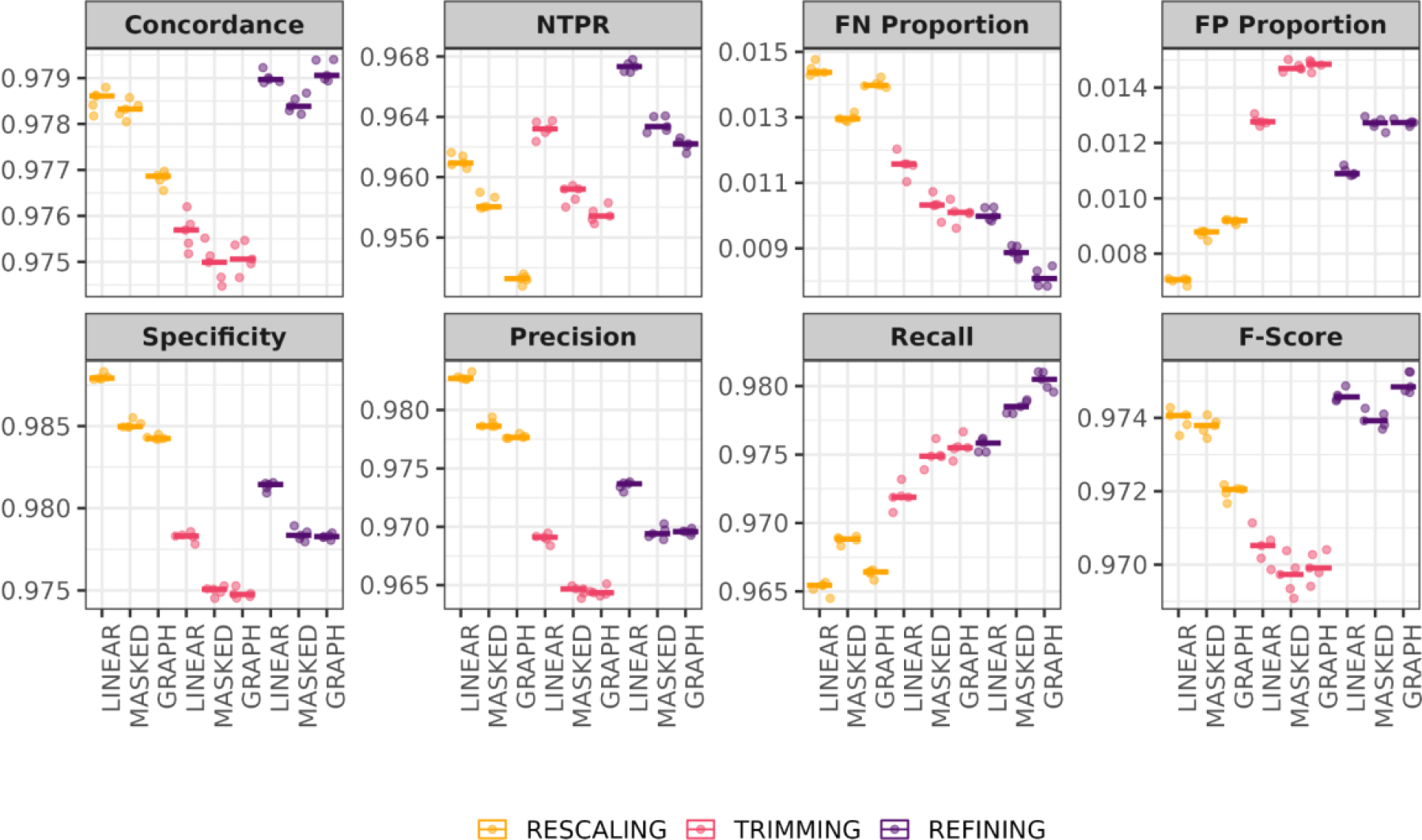
PMD-correction performances of “REFINING”, “TRIMMING”, and “REFINING” on simulated ancient genomes, calculated by comparing diploid calls with true genotypes (Figure S15).

Our results showed that the proportion of false negatives (true alternative allele called as reference) was the highest using “RESCALING”, consistent with our observations above. The NTPR (true alternative allele called as an alternative) of “RESCALING” was also the lowest.

Meanwhile, we observed <1% false positives (true reference allele called as an alternative) using “RESCALING”, compared with 1-1.5% false positives using “TRIMMING” and “REFINING”. This suggests residual PMD effects (PMD beyond the masked 10 bp) that could not be corrected using the latter two methods. These PMD-induced errors, however, were not biased towards the reference or alternative allele (Figures S2 and S9-12).

Overall, “REFINING” with *bamRefine* emerged as the top performer across the majority of evaluated indices, including concordance, recall, and F-score, suggesting it can call the largest numbers of genotypes with the least error and bias (Figure 5). The F-score values of “REFINING” were highest using the “GRAPH” alignment strategy, in contrast to “RESCALING”. Meanwhile, the “TRIMMING” strategy, which involves aggressive masking of 10 bp at the end of reads, leads to significant data loss and reduces overall depth per site (Figure S16), leading to the lowest concordance rates and F-scores in our simulations. “RESCALING” had intermediate F-scores (Figure 5) but clearly suffered from reference bias (Figures 4, Figure S10), which renders it the least useful among the three methods in our view.

Finally, we applied the two unbiased PMD-correction methods, “TRIMMING” and “REFINING”, to the 17 published ancient genomes described earlier (Table S2). When using shotgun FASTQ files and/or unfiltered BAM files and mapping using the “MASKING” or “GRAPH” strategies, neither “TRIMMING” nor “REFINING” led to reference bias (49.7%-50.7% proportion of alternative allele) (Figure 6, Table S5). However, we also noted that “TRIMMING” leads up to 2% more data loss (as measured by the number of genotyped SNPs) than “REFINING” (Figure S6, Table S6), while their error rates are comparable (Figure S17).

**Figure 6:**
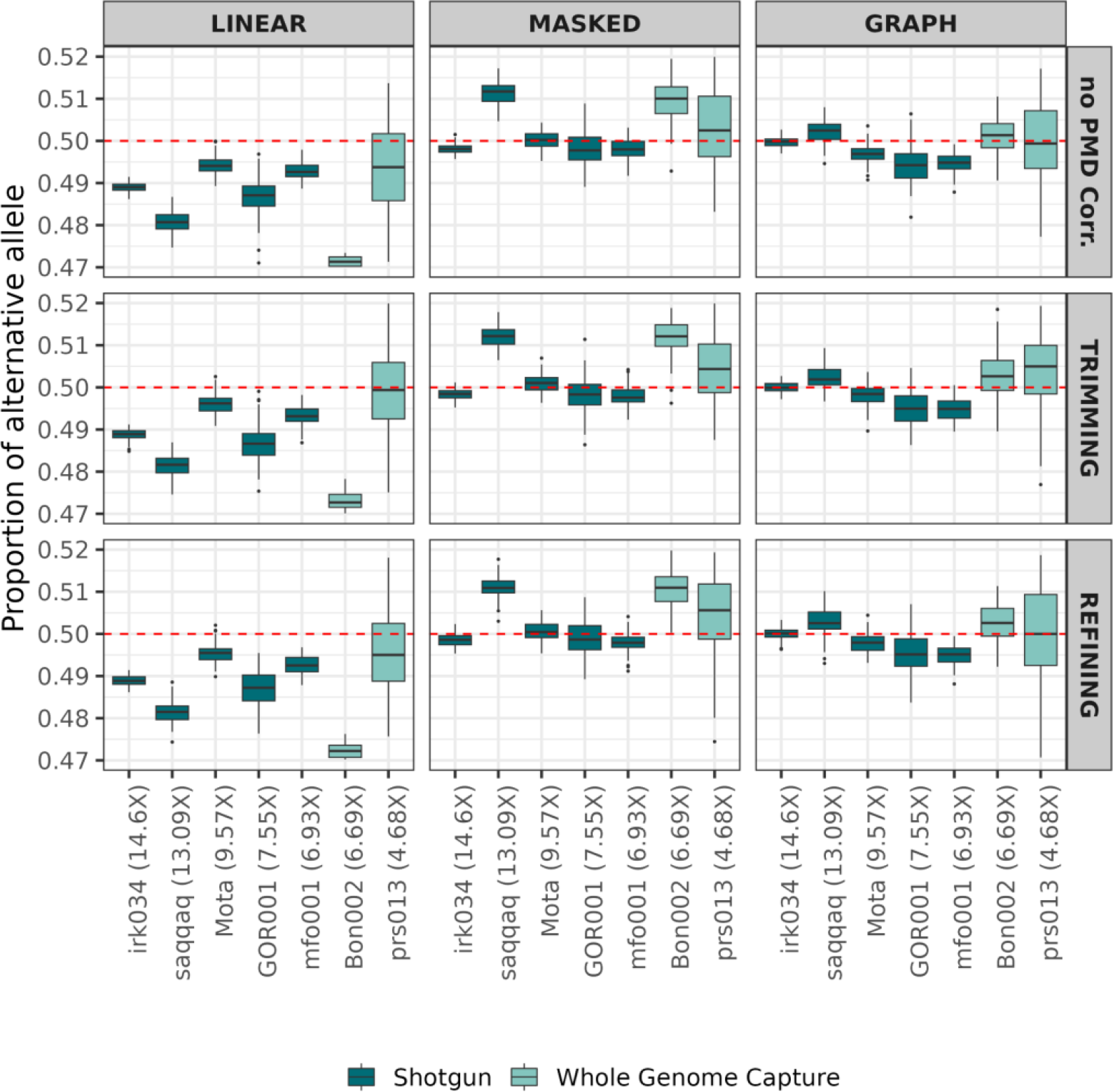
Comparing reference bias in published ancient genomes that FASTQ file available aligned with different reference genomes and PMD-effects reduced with different approaches. The plot shows the proportion of alternative alleles after randomly selecting one allele from heterozygote sites 100 times by *pileupCaller* (see also Figure S18).

### The impact of reference bias is higher on measures of allele sharing in the LINEAR strategy

Reference bias can readily lead to statistically significant asymmetries in analyses such as f_4_-statistics. We thus studied f_4_-statistics of the form f_4_(Chimp, Human Reference Genome; Ind1_MappingStrategy1, Ind1_MappingStrategy2). We found that the Human Reference Genome significantly shared more alleles with data processed using the “LINEAR” strategy (|Z| > 3) than using the “MASKED” and “GRAPH” strategies. Hence, both “MASKED” or “GRAPH” strategies largely mitigate the reference bias that arises with the “LINEAR” strategy (Figure 7A).

**Figure 7:**
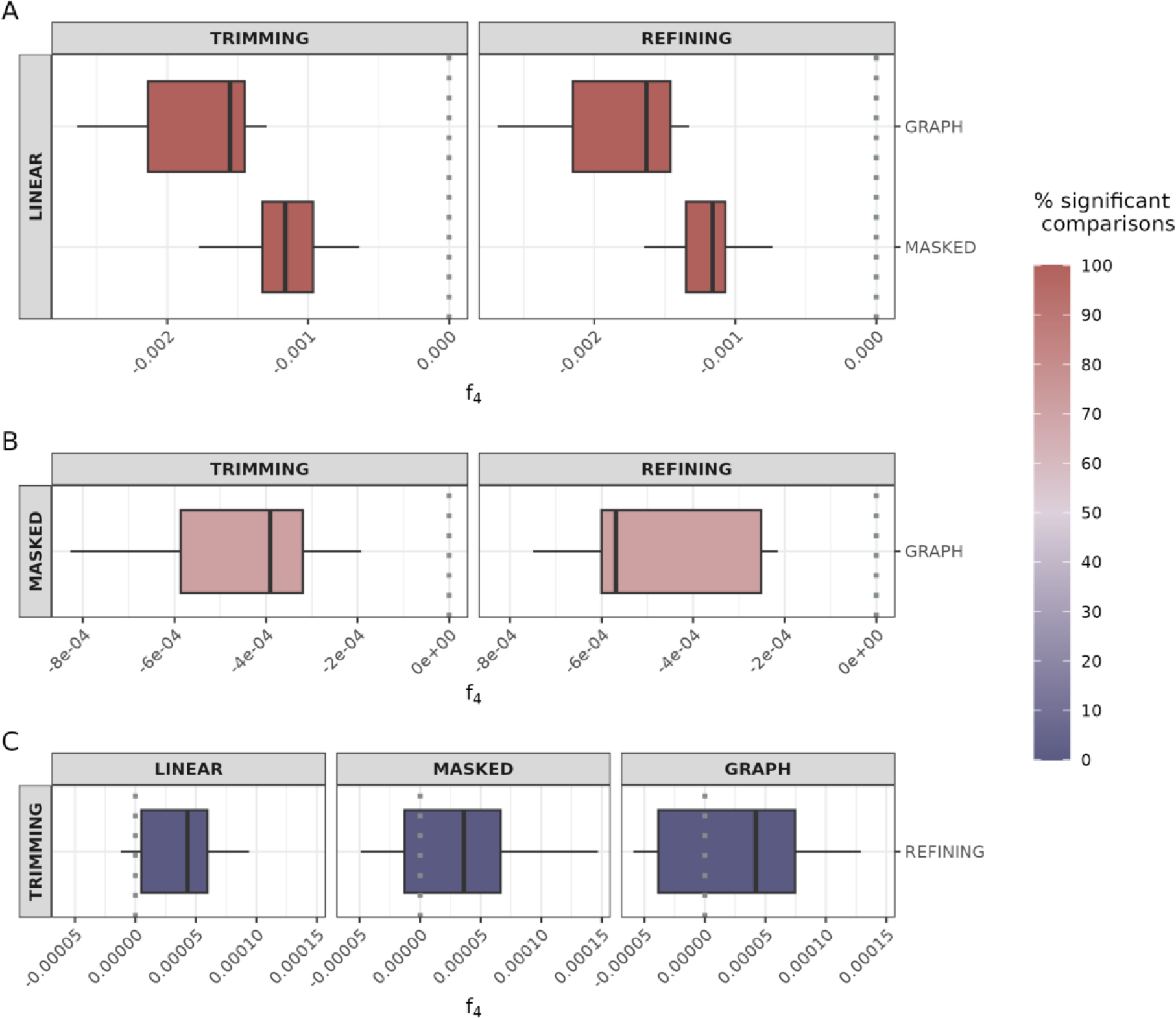
Results from the model **A)** f_4_(Chimp, Human Reference Genome; Ind1_LINEAR, Ind1_MASKED/Ind1_GRAPH), **B)** f_4_(Chimp, Human Reference Genome; Ind1_MASKED, Ind1_GRAPH) for both PMD correction strategies and **C)** f_4_(Chimp, Human Reference Genome; Ind1_Mapping_Strategy1_TRIMMING), Ind1_Mapping_Strategy1_REFINING) for all mapping strategies by using ancient genomes that FASTQ files available. The colour gradient from blue to red represents the fraction of comparisons that are nominally significant (|Z|>3) See also Figure S19 for results when the genomes with just BAM files available are used.

We further found that in 71% of comparisons, the Human Reference Genome shares more alleles with data processed using the “MASKED” strategy than the “GRAPH” strategy. This indicates that “GRAPH” is more effective in reducing reference bias, consistent with earlier results (Figure 7B).

Finally, we also compared if either “TRIMMING” or “REFINING” showed additional bias in form f_4_(Chimp, Human Reference Genome; Ind1_MappingStrategy1_TRIMMING, Ind1_MappingStrategy1_REFINING) for three mapping strategies. All results were non-significant (|Z| < 3) (Figure 7C).

## DISCUSSION

Our results confirm a strong reference bias that emerges when using the approach of linear reference genome alignment (“LINEAR”), which impacts downstream analyses such as f_4_ tests. We also find that alignment to either a masked linear reference genome (the “MASKED” strategy) or to a graph genome (“GRAPH”) effectively reduces reference bias. This observation is consistent with previous findings (5,6,8,7) and supports the feasibility of implementing these strategies for more accurate aDNA analysis. Comparing “GRAPH” and “MASKED”, the “GRAPH” approach allows a higher fraction of reads to be mapped, has higher F-scores, and appears even less affected by reference bias in f_4_ tests. Meanwhile, the “MASKED” strategy has the advantage of being simpler to implement with the standard *bwa aln* tool.

Despite their effectiveness on FASTQ data, neither “MASKED” nor “GRAPH” strategies can alleviate reference bias on paleogenome data published after mapping quality filtering. This outcome emphasizes the need for sharing all the raw data, such as the BAM files including all reads (including those with very low mapping quality), or even better, the raw FASTQ files (as mapping to a specific reference can itself create a bias). Sharing full data allows long-term healthy reusability of the data, avoiding possible batch effects due to data processing.

Meanwhile, the reference bias in SNP-capture data appears inherent to the previously widely used Agilent 1240K platform (52) and also cannot be corrected. Although Rohland and colleagues suggest that the TWIST platform is free of reference bias, this observation points to the risks introduced by experimental manipulation of ancient molecules. Imputation methods may partly help overcome such inherent biases (55), but imputation using modern-day haplotypes from specific populations may itself create new issues as, for instance, variants not present in present-day populations cannot be imputed. Overall, we believe the safest way forward for the community involves shotgun sequencing and full data sharing.

This can also allow new uses of paleogenomic data, such as copy number variation (56) or metagenomic analyses (57).

This study also introduced a new algorithm, ***bamRefine,*** for effective PMD-correction on non-UDG-treated libraries. “REFINING” with *bamRefine* selectively masks only PMD-sensitive sites at read ends and makes a larger amount of genetic information available for genotyping than the standard “TRIMMING” approach. Indeed, “REFINING” showed clearly higher performance compared to “TRIMMING” in terms of overall accuracy. “REFINING” also did not show significant reference bias, a deficiency that “RESCALING” with *mapDamage* was found to suffer from. In simulated and real datasets, the combination of “GRAPH” mapping and “REFINING” yielded the best results. Meanwhile, optimizing the “RESCALING” approach could be a worthwhile avenue for future work as it involves the lowest data loss. We also note that using UDG-treatment of aDNA is an alternative experimental solution used by a large number of laboratories.

Overall, these approaches offer promising solutions to overcome the challenges associated with aDNA analysis, extracting more information from the available data and enhancing our ability to reconstruct the population history of past populations.

## Data Availability

All samples of FASTQ/BAM files were downloaded from the accession numbers provided in the published article, except for Bon002, prs013, mfo001 and irk034, for which raw FASTQ files were obtained from the corresponding authors of the relevant publications (see Table S2).

## Acknowledgements

We thank all colleagues at the METU CompEvo and Hacettepe Human_G groups, and Torsten Günther and Anders Götherström for their support and helpful suggestions. We thank Anders Götherström, Mattias Jakobsson and Carolina Bernhardsson for sharing the raw FASTQ files for the Bon002, prs013, mfo001 and irk034 genomes. We also thank Samuel Neuenschwander for implementing *bamRefine* in the *Mapache* pipeline.

