## Supplemental Data for "Pre-processing of paleogenomes: Mitigating reference bias and postmortem damage in ancient genome data"

### Supplementary Data

This file contains Table S1-7 and Figures S1-19 to support the methodology and the main results.

**Table S1:** Number and characteristics of variants used to generate aDNA-like simulated genomes. The variants were chosen from a bi-allelic SNP set discovered in the individual 06A010111 of the Turkish Genome Project dataset (see Methods).

| | **Homozygote Reference (0/0)** | **Heterozygous (0/1)** | **Homozygore Alternative (1/1)** |
|---|---|---|---|
| **Transition** | 123,521 | 52,427 | 36,311 |
| **Transversion** | 58,994 | 25,414 | 17,080 |
| **Total** | 182,515 | 77,841 | 53,391 |

**Table S2:** Information on the published real ancient genomes used in this study. "Sample ID": the genome ID used in the relevant publication. "Data type": whether the genome was shotgun-sequenced, or sequenced after whole-genome capture or 1240K SNP capture. "Cov.": genome-wide coverage for the samples generated by shotgun or whole-genome capture and coverage of targeted 1240K SNPs for the samples generated by 1240K capture. "Location": the country where the ancient genome material was retrieved. "Input file type": the published type of data available for analysis, i.e. FASTQ, filtered BAM, or unfiltered BAM. "PMD corr. bp": how many bases at the end of the reads to PMD correction (rescaling, trimming). "Number of heterozygote SNPs": how many heterozygote SNPs are defined for each sample to use to calculate reference bias for Figures 1B, C and D. These heterozygous positions were defined as positions where alternative allele frequencies were 25-75% and had a minimum of 10 reads.

| SampleID | Data type | Cov. | Location | Publication | Input file type | PMD corr. bp | Number of heterozygote SNPs | Accession |
|---|---|---|---|---|---|---|---|---|
| GOR001 | Shotgun | 7.55 | Turkey | Koptekin et al. 2023, CurrBio | FASTQ | 10 | 35,167 | ENA: PRJEB51705 |
| irk034 | Shotgun | 14.6 | Siberia | Kılınç et al. 2021, SciAdv | FASTQ | 2 | 422,046 | Anders Götherström, Mattias Jakobsson |
| mfo001 | Shotgun | 6.93 | South Africa | Schlebusch et al. 2017, Science | FASTQ | 2 | 138,436 | Mattias Jakobsson |
| Mota | Shotgun | 9.57 | Ethiopia | Llorente et al. 2015, Science | FASTQ | 10 | 137,567 | NCBI BioProject ID: PRJNA295861 |
| Saqqaq | Shotgun | 13.09 | Greenland | Rasmussen et al. 2010, Nature | FASTQ | 2 | 96,917 | NCBI SRA: SRA010102 |
| prs013 | Whole-genome capture | 4.68 | Spain | Sanchez-Quinto et al. 2019, PNAS | FASTQ | 10 | 4,658 | Mattias Jakobsson |
| Bon002 | Whole-genome capture | 6.69 | Turkey | Kılınç et al. 2016, CurrBio | FASTQ | 10 | 29,562 | Anders Götherström, Mehmet Somel |
| LBK | Shotgun | 14.27 | Germany | Lazaridis et al. 2014, Nature | BAM (unfiltered) | 2 | 411,966 | ENA: PRJEB6272 |

| | | | | | | | | |
|---|---|---|---|---|---|---|---|---|
| UstIshim | Shotgun | 22.96 | Siberia | Fu et al. 2014, Nature | BAM (unfiltered) | 2 | 543,495 | ENA: PRJEB6622 |
| Anzick.1 | Shotgun | 11.91 | USA | Rasmussen et al. 2014, Nature | BAM (filtered) | 10 | 167,847 | http://www.cbs.dtu.dk/suppl/clovis/ |
| BOT2016 | Shotgun | 12.08 | Kazakhstan | Damgaard et al. 2018, Science | BAM (filtered) | 10 | 197,868 | ENA: PRJEB26349 |
| Yamnaya | Shotgun | 20.29 | Kazakhstan | Damgaard et al. 2018, Science | BAM (filtered) | 10 | 419,303 | ENA: PRJEB26349 |
| I0018 | 1240K Capture | 10.45 | Germany | Lipson et al. 2017, Nature | BAM (unfiltered) | 2 | 10,366 | ENA: PRJEB22629 |
| I0585 | 1240K Capture | 19.11 | Spain | Mathieson et al. 2015, Nature | BAM (unfiltered) | 2 | 19,394 | ENA: PRJEB11450 |
| I0709 | 1240K Capture | 10.65 | Turkey | Mathieson et al 2015, Nature | BAM (unfiltered) | 2 | 17,337 | ENA: PRJEB11450 |
| I2452 | 1240K Capture | 8.74 | England | Olalde et al 2018, Nature | BAM (unfiltered) | 2 | 13,042 | ENA: PRJEB23635 |
| I4106 | 1240K Capture | 5.7 | Vanuatu | Lipson et al. 2018, CurrBio | BAM (unfiltered) | 2 | 2,934 | ENA: PRJEB24938 |

**Table S3:** The proportion of alternative alleles at heterozygous sites in aDNA-like simulated genome data. The proportions were calculated by randomly selecting one allele from 77,841 heterozygote sites in aDNA-like simulated genomes (n=5 for each coverage) 100 times using pileupCaller. The “Alternative allele proportion” columns show the mean, minimum (min) and maximum (max) of the distribution of these proportions. “Coverage” indicates the depth-of-coverage. The “Alignment strategy” and “PMD correction strategy” columns indicate the methods used for alignment and PMD correction, respectively.

| **Coverage** | **Alignment strategy** | **PMD correction strategy** | **Alternative allele proportion** | | |
|---|---|---|---|---|---|
| | | | **Mean** | **Min** | **Max** |
| 0.05X | LINEAR | no PMD correction | 49.25 | 48.42 | 50.48 |
| 0.05X | LINEAR | RESCALING | 46.62 | 45.81 | 47.92 |
| 0.05X | LINEAR | TRIMMING | 49.37 | 48.02 | 51.31 |
| 0.05X | LINEAR | REFINING | 49.43 | 48.4 | 51.08 |
| 0.05X | MASKED | no PMD correction | 50.25 | 49.36 | 51.33 |
| 0.05X | MASKED | RESCALING | 47.6 | 46.75 | 48.86 |
| 0.05X | MASKED | TRIMMING | 50.25 | 48.76 | 51.96 |
| 0.05X | MASKED | REFINING | 50.36 | 49.31 | 51.83 |
| 0.05X | GRAPH | no PMD correction | 49.87 | 49.06 | 51.01 |
| 0.05X | GRAPH | RESCALING | 47.21 | 46.47 | 48.57 |
| 0.05X | GRAPH | TRIMMING | 49.95 | 48.72 | 51.64 |
| 0.05X | GRAPH | REFINING | 50.1 | 49.22 | 51.76 |
| 0.1X | LINEAR | no PMD correction | 49.27 | 48.42 | 49.97 |
| 0.1X | LINEAR | RESCALING | 46.68 | 45.81 | 47.57 |
| 0.1X | LINEAR | TRIMMING | 49.26 | 48.13 | 50.25 |
| 0.1X | LINEAR | REFINING | 49.24 | 48.29 | 50.26 |
| 0.1X | MASKED | no PMD correction | 50.33 | 49.55 | 51.01 |

| | | | | | |
|---|---|---|---|---|---|
| 0.1X | MASKED | RESCALING | 47.72 | 46.94 | 48.68 |
| 0.1X | MASKED | TRIMMING | 50.22 | 48.92 | 51.31 |
| 0.1X | MASKED | REFINING | 50.25 | 49.4 | 51.4 |
| 0.1X | GRAPH | no PMD correction | 50.05 | 49.23 | 50.8 |
| 0.1X | GRAPH | RESCALING | 47.45 | 46.56 | 48.34 |
| 0.1X | GRAPH | TRIMMING | 50.03 | 48.75 | 51.29 |
| 0.1X | GRAPH | REFINING | 50.02 | 49.06 | 51.26 |
| 1X | LINEAR | no PMD correction | 48.95 | 48.33 | 49.6 |
| 1X | LINEAR | RESCALING | 46.18 | 45.51 | 46.76 |
| 1X | LINEAR | TRIMMING | 49.19 | 48.68 | 49.69 |
| 1X | LINEAR | REFINING | 49.08 | 48.63 | 49.65 |
| 1X | MASKED | no PMD correction | 50.08 | 49.46 | 50.81 |
| 1X | MASKED | RESCALING | 47.29 | 46.61 | 47.89 |
| 1X | MASKED | TRIMMING | 50.2 | 49.71 | 50.77 |
| 1X | MASKED | REFINING | 50.17 | 49.73 | 50.65 |
| 1X | GRAPH | no PMD correction | 49.78 | 49.17 | 50.43 |
| 1X | GRAPH | RESCALING | 46.97 | 46.33 | 47.59 |
| 1X | GRAPH | TRIMMING | 49.98 | 49.55 | 50.59 |
| 1X | GRAPH | REFINING | 49.92 | 49.35 | 50.51 |
| 5X | LINEAR | no PMD correction | 48.87 | 48.29 | 49.4 |
| 5X | LINEAR | RESCALING | 45.97 | 45.44 | 46.52 |
| 5X | LINEAR | TRIMMING | 49 | 48.38 | 49.53 |
| 5X | LINEAR | REFINING | 48.95 | 48.3 | 49.46 |
| 5X | MASKED | no PMD correction | 50.22 | 49.59 | 50.75 |
| 5X | MASKED | RESCALING | 47.31 | 46.83 | 47.78 |

| | | | | | |
|---|---|---|---|---|---|
| 5X | MASKED | TRIMMING | 50.21 | 49.53 | 50.76 |
| 5X | MASKED | REFINING | 50.25 | 49.77 | 50.74 |
| 5X | GRAPH | no PMD correction | 49.87 | 49.38 | 50.45 |
| 5X | GRAPH | RESCALING | 46.93 | 46.37 | 47.51 |
| 5X | GRAPH | TRIMMING | 49.96 | 49.38 | 50.54 |
| 5X | GRAPH | REFINING | 49.93 | 49.34 | 50.45 |
| 10X | LINEAR | no PMD correction | 48.74 | 48.21 | 49.35 |
| 10X | LINEAR | RESCALING | 45.84 | 45.26 | 46.37 |
| 10X | LINEAR | TRIMMING | 48.87 | 48.48 | 49.43 |
| 10X | LINEAR | REFINING | 48.79 | 48.23 | 49.21 |
| 10X | MASKED | no PMD correction | 50.19 | 49.64 | 50.65 |
| 10X | MASKED | RESCALING | 47.26 | 46.68 | 47.76 |
| 10X | MASKED | TRIMMING | 50.19 | 49.5 | 50.74 |
| 10X | MASKED | REFINING | 50.19 | 49.66 | 50.64 |
| 10X | GRAPH | no PMD correction | 49.81 | 49.28 | 50.37 |
| 10X | GRAPH | RESCALING | 46.86 | 46.4 | 47.4 |
| 10X | GRAPH | TRIMMING | 49.89 | 49.37 | 50.46 |
| 10X | GRAPH | REFINING | 49.86 | 49.44 | 50.3 |

**Table S4:** Average number of SNPs genotyped in the aDNA-like simulated genome data. The columns “Number of SNPs” and “Number of heterozygote SNPs” show the average number of SNPs genotyped and the average number of heterozygote SNPs (used in Figure 1A) across 100 replicates. “Coverage” indicates the depth-of-coverage of chromosome 1. “Alignment strategy” and “PMD correction strategy” columns indicate the methods used for alignment and PMD correction, respectively. SE: standard error.

| Coverage | Alignment strategy | PMD correction strategy | Number of SNPs (average) | SE | Number of heterozygote SNPs (average) | SE |
|---|---|---|---|---|---|---|
| 0.05X | LINEAR | no PMD correction | 18,789 | 94 | 4,729 | 30 |
| 0.1X | LINEAR | no PMD correction | 36,479 | 136 | 9,022 | 5 |
| 1X | LINEAR | no PMD correction | 220,628 | 127 | 54,583 | 89 |
| 5X | LINEAR | no PMD correction | 310,646 | 7 | 77,035 | 7 |
| 10X | LINEAR | no PMD correction | 312,960 | 4 | 77,621 | 6 |
| 0.05X | MASKED | no PMD correction | 18,433 | 86 | 4,673 | 26 |
| 0.1X | MASKED | no PMD correction | 35,820 | 117 | 8,933 | 9 |
| 1X | MASKED | no PMD correction | 217,620 | 122 | 54,122 | 84 |
| 5X | MASKED | no PMD correction | 309,214 | 8 | 76,814 | 9 |
| 10X | MASKED | no PMD correction | 312,097 | 13 | 77,494 | 5 |
| 0.05X | GRAPH | no PMD correction | 18,775 | 95 | 4,744 | 28 |
| 0.1X | GRAPH | no PMD correction | 36,502 | 128 | 9,062 | 4 |
| 1X | GRAPH | no PMD correction | 220,996 | 153 | 54,839 | 84 |
| 5X | GRAPH | no PMD correction | 311,129 | 13 | 77,222 | 7 |
| 10X | GRAPH | no PMD correction | 313,248 | 10 | 77,711 | 2 |
| 0.05X | LINEAR | RESCALING | 18,240 | 97 | 4,502 | 29 |
| 0.1X | LINEAR | RESCALING | 35,458 | 154 | 8,607 | 9 |
| 1X | LINEAR | RESCALING | 216,851 | 128 | 53,069 | 81 |
| 5X | LINEAR | RESCALING | 310,175 | 12 | 76,892 | 8 |
| 10X | LINEAR | RESCALING | 312,896 | 4 | 77,604 | 5 |
| 0.05X | MASKED | RESCALING | 17,878 | 89 | 4,442 | 25 |

| | | | | | | |
|---|---|---|---|---|---|---|
| 0.1X | MASKED | RESCALING | 34,787 | 135 | 8,514 | 10 |
| 1X | MASKED | RESCALING | 213,768 | 127 | 52,562 | 76 |
| 5X | MASKED | RESCALING | 308,703 | 11 | 76,644 | 10 |
| 10X | MASKED | RESCALING | 312,016 | 14 | 77,463 | 5 |
| 0.05X | GRAPH | RESCALING | 18,217 | 98 | 4,510 | 27 |
| 0.1X | GRAPH | RESCALING | 35,456 | 148 | 8,638 | 14 |
| 1X | GRAPH | RESCALING | 217,115 | 157 | 53,263 | 80 |
| 5X | GRAPH | RESCALING | 310,687 | 11 | 77,081 | 6 |
| 10X | GRAPH | RESCALING | 313,201 | 9 | 77,694 | 2 |
| 0.05X | LINEAR | TRIMMING | 13,383 | 70 | 3,373 | 30 |
| 0.1X | LINEAR | TRIMMING | 26,234 | 146 | 6,490 | 15 |
| 1X | LINEAR | TRIMMING | 181,171 | 151 | 44,737 | 69 |
| 5X | LINEAR | TRIMMING | 305,346 | 19 | 75,699 | 19 |
| 10X | LINEAR | TRIMMING | 312,334 | 12 | 77,467 | 7 |
| 0.05X | MASKED | TRIMMING | 13,151 | 69 | 3,339 | 27 |
| 0.1X | MASKED | TRIMMING | 25,784 | 133 | 6,431 | 14 |
| 1X | MASKED | TRIMMING | 178,581 | 152 | 44,343 | 61 |
| 5X | MASKED | TRIMMING | 303,507 | 19 | 75,420 | 25 |
| 10X | MASKED | TRIMMING | 311,255 | 9 | 77,304 | 8 |
| 0.05X | GRAPH | TRIMMING | 13,444 | 73 | 3,403 | 30 |
| 0.1X | GRAPH | TRIMMING | 26,374 | 138 | 6,558 | 22 |
| 1X | GRAPH | TRIMMING | 181,962 | 182 | 45,090 | 63 |
| 5X | GRAPH | TRIMMING | 306,030 | 13 | 75,987 | 18 |
| 10X | GRAPH | TRIMMING | 312,704 | 10 | 77,604 | 5 |
| 0.05X | LINEAR | REFINING | 16,115 | 76 | 4,051 | 28 |
| 0.1X | LINEAR | REFINING | 31,370 | 142 | 7,756 | 9 |
| 1X | LINEAR | REFINING | 202,326 | 145 | 50,034 | 75 |
| 5X | LINEAR | REFINING | 308,703 | 7 | 76,544 | 17 |

| | | | | | | |
|---|---|---|---|---|---|---|
| 10X | LINEAR | REFINING | 312,686 | 9 | 77,557 | 6 |
| 0.05X | MASKED | REFINING | 15,817 | 72 | 4,007 | 23 |
| 0.1X | MASKED | REFINING | 30,815 | 127 | 7,682 | 7 |
| 1X | MASKED | REFINING | 199,473 | 142 | 49,595 | 75 |
| 5X | MASKED | REFINING | 307,084 | 13 | 76,297 | 19 |
| 10X | MASKED | REFINING | 311,713 | 18 | 77,411 | 8 |
| 0.05X | GRAPH | REFINING | 16,140 | 77 | 4,078 | 26 |
| 0.1X | GRAPH | REFINING | 31,457 | 133 | 7,810 | 13 |
| 1X | GRAPH | REFINING | 202,865 | 182 | 50,330 | 73 |
| 5X | GRAPH | REFINING | 309,274 | 21 | 76,777 | 15 |
| 10X | GRAPH | REFINING | 313,010 | 7 | 77,666 | 2 |

**Table S5:** The proportion of alternative alleles in real ancient genomes. We calculate the proportions after randomly selecting one allele from defined heterozygote sites (Table S4) 100 times using pileupCaller for published ancient data under different mapping and PMD correction strategies. “Sample ID”: the genome ID used in the relevant publication. “Data type”: whether the genome was shotgun-sequenced, or sequenced after whole-genome capture (“WGC”) or 1240K SNP capture. “Coverage”: genome-wide coverage for the samples generated by shotgun or whole-genome capture and coverage of targeted 1240K SNPs for the samples generated by 1240K capture. “Data type”: shotgun-sequenced or sequenced after 1240K capture. “Input file type” shows the published type of data available for analysis (FASTQ, filtered BAM, or unfiltered BAM). The “Alignment strategy” and “PMD correction strategy” columns indicate the methods used for alignment and PMD correction, respectively. “Alternative allele proportion” columns show the mean, minimum (min) and maximum (max) of the distribution of these proportions.

| **Sample ID** | **Alignment strategy** | **Coverage** | **PMD correction strategy** | **Input file** | **Data type** | **Alternative allele proportion** | | |
|---|---|---|---|---|---|---|---|---|
| | | | | | | **Mean** | **Min** | **Max** |
| irk034 | LINEAR | 14.60 | no PMD Corr. | FASTQ | Shotgun | 48.9 | 48.62 | 49.15 |
| irk034 | LINEAR | 14.60 | TRIMMING | FASTQ | Shotgun | 48.88 | 48.48 | 49.12 |
| irk034 | LINEAR | 14.60 | REFINING | FASTQ | Shotgun | 48.89 | 48.62 | 49.14 |
| irk034 | MASKED | 14.60 | no PMD Corr. | FASTQ | Shotgun | 49.82 | 49.57 | 50.15 |
| irk034 | MASKED | 14.60 | TRIMMING | FASTQ | Shotgun | 49.84 | 49.52 | 50.12 |
| irk034 | MASKED | 14.60 | REFINING | FASTQ | Shotgun | 49.85 | 49.53 | 50.24 |
| irk034 | GRAPH | 14.60 | no PMD Corr. | FASTQ | Shotgun | 49.98 | 49.7 | 50.27 |
| irk034 | GRAPH | 14.60 | TRIMMING | FASTQ | Shotgun | 49.99 | 49.72 | 50.27 |
| irk034 | GRAPH | 14.60 | REFINING | FASTQ | Shotgun | 50.01 | 49.63 | 50.33 |
| saqqaq | LINEAR | 13.09 | no PMD Corr. | FASTQ | Shotgun | 48.08 | 47.47 | 48.67 |
| saqqaq | LINEAR | 13.09 | TRIMMING | FASTQ | Shotgun | 48.14 | 47.46 | 48.7 |
| saqqaq | LINEAR | 13.09 | REFINING | FASTQ | Shotgun | 48.15 | 47.43 | 48.85 |
| saqqaq | MASKED | 13.09 | no PMD Corr. | FASTQ | Shotgun | 51.13 | 50.47 | 51.72 |
| saqqaq | MASKED | 13.09 | TRIMMING | FASTQ | Shotgun | 51.2 | 50.65 | 51.79 |
| saqqaq | MASKED | 13.09 | REFINING | FASTQ | Shotgun | 51.1 | 50.3 | 51.77 |

| | | | | | | | | |
|---|---|---|---|---|---|---|---|---|
| saqqaq | GRAPH | 13.09 | no PMD Corr. | FASTQ | Shotgun | 50.21 | 49.46 | 50.8 |
| saqqaq | GRAPH | 13.09 | TRIMMING | FASTQ | Shotgun | 50.23 | 49.67 | 50.93 |
| saqqaq | GRAPH | 13.09 | REFINING | FASTQ | Shotgun | 50.27 | 49.3 | 51.02 |
| Mota | LINEAR | 9.57 | no PMD Corr. | FASTQ | Shotgun | 49.42 | 48.93 | 49.99 |
| Mota | LINEAR | 9.57 | TRIMMING | FASTQ | Shotgun | 49.6 | 49.09 | 50.26 |
| Mota | LINEAR | 9.57 | REFINING | FASTQ | Shotgun | 49.54 | 48.99 | 50.21 |
| Mota | MASKED | 9.57 | no PMD Corr. | FASTQ | Shotgun | 50.02 | 49.52 | 50.43 |
| Mota | MASKED | 9.57 | TRIMMING | FASTQ | Shotgun | 50.1 | 49.63 | 50.69 |
| Mota | MASKED | 9.57 | REFINING | FASTQ | Shotgun | 50.05 | 49.54 | 50.56 |
| Mota | GRAPH | 9.57 | no PMD Corr. | FASTQ | Shotgun | 49.71 | 49.08 | 50.35 |
| Mota | GRAPH | 9.57 | TRIMMING | FASTQ | Shotgun | 49.82 | 48.97 | 50.37 |
| Mota | GRAPH | 9.57 | REFINING | FASTQ | Shotgun | 49.77 | 49.31 | 50.44 |
| GOR001 | LINEAR | 7.55 | no PMD Corr. | FASTQ | Shotgun | 48.67 | 47.1 | 49.69 |
| GOR001 | LINEAR | 7.55 | TRIMMING | FASTQ | Shotgun | 48.67 | 47.54 | 49.91 |
| GOR001 | LINEAR | 7.55 | REFINING | FASTQ | Shotgun | 48.73 | 47.63 | 49.55 |
| GOR001 | MASKED | 7.55 | no PMD Corr. | FASTQ | Shotgun | 49.79 | 48.91 | 50.89 |
| GOR001 | MASKED | 7.55 | TRIMMING | FASTQ | Shotgun | 49.83 | 48.64 | 51.14 |
| GOR001 | MASKED | 7.55 | REFINING | FASTQ | Shotgun | 49.89 | 48.93 | 50.87 |
| GOR001 | GRAPH | 7.55 | no PMD Corr. | FASTQ | Shotgun | 49.42 | 48.19 | 50.64 |
| GOR001 | GRAPH | 7.55 | TRIMMING | FASTQ | Shotgun | 49.49 | 48.63 | 50.46 |
| GOR001 | GRAPH | 7.55 | REFINING | FASTQ | Shotgun | 49.56 | 48.37 | 50.71 |
| mfo001 | LINEAR | 6.93 | no PMD Corr. | FASTQ | Shotgun | 49.29 | 48.87 | 49.79 |
| mfo001 | LINEAR | 6.93 | TRIMMING | FASTQ | Shotgun | 49.32 | 48.69 | 49.82 |
| mfo001 | LINEAR | 6.93 | REFINING | FASTQ | Shotgun | 49.27 | 48.79 | 49.68 |
| mfo001 | MASKED | 6.93 | no PMD Corr. | FASTQ | Shotgun | 49.8 | 49.17 | 50.31 |

| | | | | | | | | |
|---|---|---|---|---|---|---|---|---|
| mfo001 | MASKED | 6.93 | TRIMMING | FASTQ | Shotgun | 49.79 | 49.23 | 50.42 |
| mfo001 | MASKED | 6.93 | REFINING | FASTQ | Shotgun | 49.79 | 49.12 | 50.41 |
| mfo001 | GRAPH | 6.93 | no PMD Corr. | FASTQ | Shotgun | 49.48 | 48.79 | 49.92 |
| mfo001 | GRAPH | 6.93 | TRIMMING | FASTQ | Shotgun | 49.48 | 48.95 | 50.05 |
| mfo001 | GRAPH | 6.93 | REFINING | FASTQ | Shotgun | 49.5 | 48.81 | 49.95 |
| Bon002 | LINEAR | 6.69 | no PMD Corr. | FASTQ | WGC | 46.4 | 45.1 | 47.34 |
| Bon002 | LINEAR | 6.69 | TRIMMING | FASTQ | WGC | 46.96 | 45.84 | 47.83 |
| Bon002 | LINEAR | 6.69 | REFINING | FASTQ | WGC | 46.61 | 45.41 | 47.63 |
| Bon002 | MASKED | 6.69 | no PMD Corr. | FASTQ | WGC | 51.02 | 49.28 | 52.5 |
| Bon002 | MASKED | 6.69 | TRIMMING | FASTQ | WGC | 51.28 | 49.62 | 52.79 |
| Bon002 | MASKED | 6.69 | REFINING | FASTQ | WGC | 51.06 | 50.03 | 51.98 |
| Bon002 | GRAPH | 6.69 | no PMD Corr. | FASTQ | WGC | 50.12 | 49.06 | 51.05 |
| Bon002 | GRAPH | 6.69 | TRIMMING | FASTQ | WGC | 50.29 | 48.96 | 51.85 |
| Bon002 | GRAPH | 6.69 | REFINING | FASTQ | WGC | 50.27 | 49.22 | 51.13 |
| prs013 | LINEAR | 4.68 | no PMD Corr. | FASTQ | WGC | 49.32 | 46.2 | 53.12 |
| prs013 | LINEAR | 4.68 | TRIMMING | FASTQ | WGC | 49.85 | 47.51 | 52 |
| prs013 | LINEAR | 4.68 | REFINING | FASTQ | WGC | 49.63 | 46.63 | 53.93 |
| prs013 | MASKED | 4.68 | no PMD Corr. | FASTQ | WGC | 50.66 | 48.32 | 54.24 |
| prs013 | MASKED | 4.68 | TRIMMING | FASTQ | WGC | 50.92 | 48.75 | 54.86 |
| prs013 | MASKED | 4.68 | REFINING | FASTQ | WGC | 50.64 | 47.44 | 53.12 |
| prs013 | GRAPH | 4.68 | no PMD Corr. | FASTQ | WGC | 50.12 | 47.73 | 53.15 |
| prs013 | GRAPH | 4.68 | TRIMMING | FASTQ | WGC | 50.48 | 46.63 | 52.81 |
| prs013 | GRAPH | 4.68 | REFINING | FASTQ | WGC | 50.25 | 47.07 | 53.55 |
| UstIshim | LINEAR | 22.96 | no PMD Corr. | BAM | Shotgun (unfiltered) | 49.09 | 48.82 | 49.37 |
| UstIshim | LINEAR | 22.96 | TRIMMING | BAM | Shotgun (unfiltered) | 49.14 | 48.86 | 49.38 |

| | | | | | | | | |
|---|---|---|---|---|---|---|---|---|
| UstIshim | LINEAR | 22.96 | REFINING | BAM | Shotgun (unfiltered) | 49.11 | 48.88 | 49.38 |
| UstIshim | MASKED | 22.96 | no PMD Corr. | BAM | Shotgun (unfiltered) | 50.06 | 49.8 | 50.31 |
| UstIshim | MASKED | 22.96 | TRIMMING | BAM | Shotgun (unfiltered) | 50.07 | 49.77 | 50.32 |
| UstIshim | MASKED | 22.96 | REFINING | BAM | Shotgun (unfiltered) | 50.07 | 49.89 | 50.31 |
| UstIshim | GRAPH | 22.96 | no PMD Corr. | BAM | Shotgun (unfiltered) | 49.77 | 49.49 | 50.02 |
| UstIshim | GRAPH | 22.96 | TRIMMING | BAM | Shotgun (unfiltered) | 49.79 | 49.52 | 50.08 |
| UstIshim | GRAPH | 22.96 | REFINING | BAM | Shotgun (unfiltered) | 49.77 | 49.53 | 49.98 |
| LBK | LINEAR | 14.27 | no PMD Corr. | BAM | Shotgun (unfiltered) | 49.43 | 49.12 | 49.77 |
| LBK | LINEAR | 14.27 | TRIMMING | BAM | Shotgun (unfiltered) | 49.49 | 49.07 | 49.77 |
| LBK | LINEAR | 14.27 | REFINING | BAM | Shotgun (unfiltered) | 49.43 | 48.99 | 49.72 |
| LBK | MASKED | 14.27 | no PMD Corr. | BAM | Shotgun (unfiltered) | 50.1 | 49.8 | 50.42 |
| LBK | MASKED | 14.27 | TRIMMING | BAM | Shotgun (unfiltered) | 50.14 | 49.75 | 50.49 |
| LBK | MASKED | 14.27 | REFINING | BAM | Shotgun (unfiltered) | 50.14 | 49.79 | 50.47 |
| LBK | GRAPH | 14.27 | no PMD Corr. | BAM | Shotgun (unfiltered) | 49.85 | 49.56 | 50.31 |
| LBK | GRAPH | 14.27 | TRIMMING | BAM | Shotgun (unfiltered) | 49.86 | 49.58 | 50.13 |
| LBK | GRAPH | 14.27 | REFINING | BAM | Shotgun (unfiltered) | 49.85 | 49.56 | 50.12 |
| Yamnaya | LINEAR | 20.29 | no PMD Corr. | BAM | Shotgun (filtered) | 48.09 | 47.72 | 48.42 |
| Yamnaya | LINEAR | 20.29 | TRIMMING | BAM | Shotgun (filtered) | 48.29 | 48.01 | 48.57 |
| Yamnaya | LINEAR | 20.29 | REFINING | BAM | Shotgun (filtered) | 48.17 | 47.85 | 48.52 |
| Yamnaya | MASKED | 20.29 | no PMD Corr. | BAM | Shotgun (filtered) | 48.88 | 48.5 | 49.34 |
| Yamnaya | MASKED | 20.29 | TRIMMING | BAM | Shotgun (filtered) | 49.04 | 48.71 | 49.4 |
| Yamnaya | MASKED | 20.29 | REFINING | BAM | Shotgun (filtered) | 48.95 | 48.55 | 49.29 |
| Yamnaya | GRAPH | 20.29 | no PMD Corr. | BAM | Shotgun (filtered) | 48.55 | 48.16 | 48.94 |
| Yamnaya | GRAPH | 20.29 | TRIMMING | BAM | Shotgun (filtered) | 48.72 | 48.44 | 49.07 |
| Yamnaya | GRAPH | 20.29 | REFINING | BAM | Shotgun (filtered) | 48.64 | 48.36 | 48.93 |

| | | | | | | | | |
|---|---|---|---|---|---|---|---|---|
| BOT2016 | LINEAR | 12.08 | no PMD Corr. | BAM | Shotgun (filtered) | 48.37 | 47.94 | 48.82 |
| BOT2016 | LINEAR | 12.08 | TRIMMING | BAM | Shotgun (filtered) | 48.54 | 47.97 | 49.08 |
| BOT2016 | LINEAR | 12.08 | REFINING | BAM | Shotgun (filtered) | 48.46 | 47.98 | 49.16 |
| BOT2016 | MASKED | 12.08 | no PMD Corr. | BAM | Shotgun (filtered) | 49.16 | 48.76 | 49.56 |
| BOT2016 | MASKED | 12.08 | TRIMMING | BAM | Shotgun (filtered) | 49.35 | 48.82 | 49.77 |
| BOT2016 | MASKED | 12.08 | REFINING | BAM | Shotgun (filtered) | 49.24 | 48.81 | 49.69 |
| BOT2016 | GRAPH | 12.08 | no PMD Corr. | BAM | Shotgun (filtered) | 48.84 | 48.28 | 49.33 |
| BOT2016 | GRAPH | 12.08 | TRIMMING | BAM | Shotgun (filtered) | 49.01 | 48.52 | 49.54 |
| BOT2016 | GRAPH | 12.08 | REFINING | BAM | Shotgun (filtered) | 48.89 | 48.4 | 49.35 |
| Anzick.1 | LINEAR | 11.91 | no PMD Corr. | BAM | Shotgun (filtered) | 47.39 | 46.86 | 47.97 |
| Anzick.1 | LINEAR | 11.91 | TRIMMING | BAM | Shotgun (filtered) | 47.63 | 47.13 | 48.21 |
| Anzick.1 | LINEAR | 11.91 | REFINING | BAM | Shotgun (filtered) | 47.56 | 47.01 | 48.01 |
| Anzick.1 | MASKED | 11.91 | no PMD Corr. | BAM | Shotgun (filtered) | 48.46 | 47.85 | 49.03 |
| Anzick.1 | MASKED | 11.91 | TRIMMING | BAM | Shotgun (filtered) | 48.65 | 48.08 | 49.12 |
| Anzick.1 | MASKED | 11.91 | REFINING | BAM | Shotgun (filtered) | 48.56 | 47.95 | 49 |
| Anzick.1 | GRAPH | 11.91 | no PMD Corr. | BAM | Shotgun (filtered) | 47.85 | 47.24 | 48.43 |
| Anzick.1 | GRAPH | 11.91 | TRIMMING | BAM | Shotgun (filtered) | 48.02 | 47.47 | 48.63 |
| Anzick.1 | GRAPH | 11.91 | REFINING | BAM | Shotgun (filtered) | 47.93 | 47.53 | 48.41 |
| I0585 | LINEAR | 19.11 | no PMD Corr. | BAM | 1240K Capture | 49.38 | 47.39 | 50.6 |
| I0585 | LINEAR | 19.11 | TRIMMING | BAM | 1240K Capture | 49.5 | 47.79 | 51.33 |
| I0585 | LINEAR | 19.11 | REFINING | BAM | 1240K Capture | 49.46 | 48.38 | 50.66 |
| I0585 | MASKED | 19.11 | no PMD Corr. | BAM | 1240K Capture | 49.52 | 47.66 | 51.05 |
| I0585 | MASKED | 19.11 | TRIMMING | BAM | 1240K Capture | 49.57 | 48.44 | 50.93 |
| I0585 | MASKED | 19.11 | REFINING | BAM | 1240K Capture | 49.52 | 48.13 | 50.78 |
| I0585 | GRAPH | 19.11 | no PMD Corr. | BAM | 1240K Capture | 49.43 | 48.2 | 50.9 |

| | | | | | | | | |
|---|---|---|---|---|---|---|---|---|
| I0585 | GRAPH | 19.11 | TRIMMING | BAM | 1240K Capture | 49.59 | 48.52 | 50.78 |
| I0585 | GRAPH | 19.11 | REFINING | BAM | 1240K Capture | 49.5 | 47.99 | 50.68 |
| I0709 | LINEAR | 10.65 | no PMD Corr. | BAM | 1240K Capture | 49.2 | 47.58 | 50.68 |
| I0709 | LINEAR | 10.65 | TRIMMING | BAM | 1240K Capture | 49.24 | 47.86 | 51.14 |
| I0709 | LINEAR | 10.65 | REFINING | BAM | 1240K Capture | 49.37 | 47.64 | 51.13 |
| I0709 | MASKED | 10.65 | no PMD Corr. | BAM | 1240K Capture | 49.3 | 47.24 | 50.7 |
| I0709 | MASKED | 10.65 | TRIMMING | BAM | 1240K Capture | 49.41 | 47.75 | 50.84 |
| I0709 | MASKED | 10.65 | REFINING | BAM | 1240K Capture | 49.45 | 47.92 | 50.92 |
| I0709 | GRAPH | 10.65 | no PMD Corr. | BAM | 1240K Capture | 49.36 | 47.94 | 51.01 |
| I0709 | GRAPH | 10.65 | TRIMMING | BAM | 1240K Capture | 49.28 | 47.76 | 50.65 |
| I0709 | GRAPH | 10.65 | REFINING | BAM | 1240K Capture | 49.18 | 47.84 | 50.98 |
| I0018 | LINEAR | 10.45 | no PMD Corr. | BAM | 1240K Capture | 48.83 | 46.94 | 50.48 |
| I0018 | LINEAR | 10.45 | TRIMMING | BAM | 1240K Capture | 49.07 | 47.25 | 50.74 |
| I0018 | LINEAR | 10.45 | REFINING | BAM | 1240K Capture | 48.94 | 46.16 | 50.82 |
| I0018 | MASKED | 10.45 | no PMD Corr. | BAM | 1240K Capture | 49.05 | 46.88 | 51 |
| I0018 | MASKED | 10.45 | TRIMMING | BAM | 1240K Capture | 48.88 | 46.38 | 50.77 |
| I0018 | MASKED | 10.45 | REFINING | BAM | 1240K Capture | 49.06 | 47.06 | 51.04 |
| I0018 | GRAPH | 10.45 | no PMD Corr. | BAM | 1240K Capture | 48.82 | 46.56 | 50.92 |
| I0018 | GRAPH | 10.45 | TRIMMING | BAM | 1240K Capture | 49.03 | 46.87 | 50.77 |
| I0018 | GRAPH | 10.45 | REFINING | BAM | 1240K Capture | 48.94 | 47.03 | 50.6 |
| I2452 | LINEAR | 8.74 | no PMD Corr. | BAM | 1240K Capture | 49.3 | 48.02 | 51.14 |
| I2452 | LINEAR | 8.74 | TRIMMING | BAM | 1240K Capture | 49.27 | 47.73 | 50.64 |
| I2452 | LINEAR | 8.74 | REFINING | BAM | 1240K Capture | 49.35 | 47.86 | 50.78 |
| I2452 | MASKED | 8.74 | no PMD Corr. | BAM | 1240K Capture | 49.29 | 47.32 | 50.73 |
| I2452 | MASKED | 8.74 | TRIMMING | BAM | 1240K Capture | 49.26 | 47.28 | 51.16 |

| | | | | | | | | |
|---|---|---|---|---|---|---|---|---|
| I2452 | MASKED | 8.74 | REFINING | BAM | 1240K Capture | 49.31 | 47.23 | 51.14 |
| I2452 | GRAPH | 8.74 | no PMD Corr. | BAM | 1240K Capture | 49.33 | 47.8 | 51.07 |
| I2452 | GRAPH | 8.74 | TRIMMING | BAM | 1240K Capture | 49.21 | 47.32 | 51.39 |
| I2452 | GRAPH | 8.74 | REFINING | BAM | 1240K Capture | 49.24 | 47.37 | 50.8 |
| I4106 | LINEAR | 5.70 | no PMD Corr. | BAM | 1240K Capture | 49.42 | 45.71 | 52.67 |
| I4106 | LINEAR | 5.70 | TRIMMING | BAM | 1240K Capture | 49.04 | 44.6 | 52.88 |
| I4106 | LINEAR | 5.70 | REFINING | BAM | 1240K Capture | 49.22 | 45.01 | 53.08 |
| I4106 | MASKED | 5.70 | no PMD Corr. | BAM | 1240K Capture | 49.84 | 43.9 | 53.58 |
| I4106 | MASKED | 5.70 | TRIMMING | BAM | 1240K Capture | 49.51 | 45.51 | 53.48 |
| I4106 | MASKED | 5.70 | REFINING | BAM | 1240K Capture | 49.76 | 46.72 | 52.98 |
| I4106 | GRAPH | 5.70 | no PMD Corr. | BAM | 1240K Capture | 49.68 | 45.51 | 52.98 |
| I4106 | GRAPH | 5.70 | TRIMMING | BAM | 1240K Capture | 49.3 | 45.91 | 52.67 |
| I4106 | GRAPH | 5.70 | REFINING | BAM | 1240K Capture | 49.41 | 44.8 | 53.08 |

**Table S6:** Number of SNPs genotyped in real ancient genomes. The column “Number of heterozygote SNPs” shows the average number of genotyped heterozygote SNPs used in Figure S1B-D. “Sample ID”: the genome ID used in the relevant publication. “Data type”: whether the genome was shotgun-sequenced, or sequenced after whole-genome capture or 1240K SNP capture. The “Coverage” column indicates genome-wide coverage for the samples generated by shotgun or whole-genome capture and coverage of targeted 1240K SNPs for the samples generated by 1240K capture. “Data type”: shotgun-sequenced or sequenced after 1240K capture. “Input file type” shows the published type of data available for analysis (FASTQ, filtered BAM, or unfiltered BAM). “Alignment strategy” and “PMD correction strategy” columns indicate the methods used for alignment and PMD correction, respectively. “Number of SNPs” and “Number of heterozygous SNPs” columns show the total number of SNPs and heterozygous SNPs called from each genome, respectively.

| **Sample ID** | **Alignment strategy** | **PMD correction strategy** | **Coverage** | **Input file type** | **Data type** | **Number of SNPs** | **Number of heterozygous SNPs** |
|---|---|---|---|---|---|---|---|
| GOR001 | LINEAR | no PMD correction | 7.55 | FASTQ | Shotgun | 4,750,429 | 12,241 |
| Mota | LINEAR | no PMD correction | 9.57 | FASTQ | Shotgun | 4,718,552 | 48,387 |
| irk034 | LINEAR | no PMD correction | 14.60 | FASTQ | Shotgun | 4,767,436 | 150,413 |
| mfo001 | LINEAR | no PMD correction | 6.93 | FASTQ | Shotgun | 4,728,593 | 48,480 |
| saqqaq | LINEAR | no PMD correction | 13.09 | FASTQ | Shotgun | 4,268,599 | 31,037 |
| Bon002 | LINEAR | no PMD correction | 6.69 | FASTQ | Whole Genome Capture | 4,508,651 | 10,405 |
| prs013 | LINEAR | no PMD correction | 4.68 | FASTQ | Whole Genome Capture | 4,632,469 | 1,604 |
| I0018 | LINEAR | no PMD correction | 10.45 | BAM | 1240K Capture | 1,075,119 | 3,643 |
| I0585 | LINEAR | no PMD correction | 19.11 | BAM | 1240K Capture | 2,204,707 | 7,018 |
| I0709 | LINEAR | no PMD correction | 10.65 | BAM | 1240K Capture | 1,373,275 | 6,028 |
| I2452 | LINEAR | no PMD correction | 8.74 | BAM | 1240K Capture | 878,294 | 4,425 |
| I4106 | LINEAR | no PMD correction | 5.70 | BAM | 1240K Capture | 865,774 | 991 |
| Anzick.1 | LINEAR | no PMD correction | 11.91 | BAM | Shotgun (filtered) | 4,658,300 | 55,642 |

| | | | | | | | |
|---|---|---|---|---|---|---|---|
| BOT2016 | LINEAR | no PMD correction | 12.08 | BAM | Shotgun (filtered) | 4,763,717 | 70,033 |
| Yamnaya | LINEAR | no PMD correction | 20.29 | BAM | Shotgun (filtered) | 4,765,267 | 149,249 |
| LBK | LINEAR | no PMD correction | 14.27 | BAM | Shotgun (unfiltered) | 4,740,206 | 144,195 |
| UstIshim | LINEAR | no PMD correction | 22.96 | BAM | Shotgun (unfiltered) | 4,749,265 | 192,946 |
| GOR001 | LINEAR | REFINING | 7.55 | FASTQ | Shotgun | 4,742,362 | 35,167 |
| Mota | LINEAR | REFINING | 9.57 | FASTQ | Shotgun | 4,707,108 | 137,567 |
| irk034 | LINEAR | REFINING | 14.60 | FASTQ | Shotgun | 4,767,220 | 422,046 |
| mfo001 | LINEAR | REFINING | 6.93 | FASTQ | Shotgun | 4,725,970 | 138,436 |
| saqqaq | LINEAR | REFINING | 13.09 | FASTQ | Shotgun | 4,248,284 | 96,917 |
| Bon002 | LINEAR | REFINING | 6.69 | FASTQ | Whole Genome Capture | 4,412,423 | 29,562 |
| prs013 | LINEAR | REFINING | 4.68 | FASTQ | Whole Genome Capture | 4,565,265 | 4,658 |
| I0018 | LINEAR | REFINING | 10.45 | BAM | 1240K Capture | 1,053,294 | 10,366 |
| I0585 | LINEAR | REFINING | 19.11 | BAM | 1240K Capture | 2,159,594 | 19,394 |
| I0709 | LINEAR | REFINING | 10.65 | BAM | 1240K Capture | 1,347,384 | 17,337 |
| I2452 | LINEAR | REFINING | 8.74 | BAM | 1240K Capture | 861,702 | 13,042 |
| I4106 | LINEAR | REFINING | 5.70 | BAM | 1240K Capture | 848,359 | 2,934 |
| Anzick.1 | LINEAR | REFINING | 11.91 | BAM | Shotgun (filtered) | 4,621,821 | 167,847 |
| BOT2016 | LINEAR | REFINING | 12.08 | BAM | Shotgun (filtered) | 4,761,310 | 197,868 |
| Yamnaya | LINEAR | REFINING | 20.29 | BAM | Shotgun (filtered) | 4,763,683 | 419,303 |
| LBK | LINEAR | REFINING | 14.27 | BAM | Shotgun (unfiltered) | 4,739,268 | 411,966 |
| UstIshim | LINEAR | REFINING | 22.96 | BAM | Shotgun (unfiltered) | 4,748,625 | 543,495 |
| GOR001 | LINEAR | TRIMMING | 7.55 | FASTQ | Shotgun | 4,729,997 | 35,167 |

| | | | | | | | |
|---|---|---|---|---|---|---|---|
| Mota | LINEAR | TRIMMING | 9.57 | FASTQ | Shotgun | 4,691,679 | 137,567 |
| irk034 | LINEAR | TRIMMING | 14.60 | FASTQ | Shotgun | 4,766,990 | 422,046 |
| mfo001 | LINEAR | TRIMMING | 6.93 | FASTQ | Shotgun | 4,723,409 | 138,436 |
| saqqaq | LINEAR | TRIMMING | 13.09 | FASTQ | Shotgun | 4,223,538 | 96,917 |
| Bon002 | LINEAR | TRIMMING | 6.69 | FASTQ | Whole Genome Capture | 4,290,794 | 29,562 |
| prs013 | LINEAR | TRIMMING | 4.68 | FASTQ | Whole Genome Capture | 4,453,850 | 4,658 |
| I0018 | LINEAR | TRIMMING | 10.45 | BAM | 1240K Capture | 1,028,984 | 10,366 |
| I0585 | LINEAR | TRIMMING | 19.11 | BAM | 1240K Capture | 2,116,481 | 19,394 |
| I0709 | LINEAR | TRIMMING | 10.65 | BAM | 1240K Capture | 1,321,811 | 17,337 |
| I2452 | LINEAR | TRIMMING | 8.74 | BAM | 1240K Capture | 845,259 | 13,042 |
| I4106 | LINEAR | TRIMMING | 5.70 | BAM | 1240K Capture | 832,026 | 2,934 |
| Anzick.1 | LINEAR | TRIMMING | 11.91 | BAM | Shotgun (filtered) | 4,568,432 | 167,847 |
| BOT2016 | LINEAR | TRIMMING | 12.08 | BAM | Shotgun (filtered) | 4,758,199 | 197,868 |
| Yamnaya | LINEAR | TRIMMING | 20.29 | BAM | Shotgun (filtered) | 4,761,784 | 419,303 |
| LBK | LINEAR | TRIMMING | 14.27 | BAM | Shotgun (unfiltered) | 4,738,100 | 411,966 |
| UstIshim | LINEAR | TRIMMING | 22.96 | BAM | Shotgun (unfiltered) | 4,747,958 | 543,495 |
| GOR001 | MASKED | no PMD correction | 7.55 | FASTQ | Shotgun | 4,730,189 | 12,241 |
| Mota | MASKED | no PMD correction | 9.57 | FASTQ | Shotgun | 4,686,258 | 48,387 |
| irk034 | MASKED | no PMD correction | 14.60 | FASTQ | Shotgun | 4,762,930 | 150,413 |
| mfo001 | MASKED | no PMD correction | 6.93 | FASTQ | Shotgun | 4,704,348 | 48,480 |
| saqqaq | MASKED | no PMD correction | 13.09 | FASTQ | Shotgun | 4,178,610 | 31,031 |
| Bon002 | MASKED | no PMD correction | 6.69 | FASTQ | Whole Genome Capture | 4,484,504 | 10,403 |

| | | | | | | | |
|---|---|---|---|---|---|---|---|
| prs013 | MASKED | no PMD correction | 4.68 | FASTQ | Whole Genome Capture | 4,586,305 | 1,604 |
| I0018 | MASKED | no PMD correction | 10.45 | BAM | 1240K Capture | 1,044,328 | 3,643 |
| I0585 | MASKED | no PMD correction | 19.11 | BAM | 1240K Capture | 2,146,722 | 7,017 |
| I0709 | MASKED | no PMD correction | 10.65 | BAM | 1240K Capture | 1,330,682 | 6,029 |
| I2452 | MASKED | no PMD correction | 8.74 | BAM | 1240K Capture | 852,092 | 4,425 |
| I4106 | MASKED | no PMD correction | 5.70 | BAM | 1240K Capture | 838,323 | 991 |
| Anzick.1 | MASKED | no PMD correction | 11.91 | BAM | Shotgun (filtered) | 4,627,071 | 55,640 |
| BOT2016 | MASKED | no PMD correction | 12.08 | BAM | Shotgun (filtered) | 4,754,010 | 70,031 |
| Yamnaya | MASKED | no PMD correction | 20.29 | BAM | Shotgun (filtered) | 4,755,398 | 149,246 |
| LBK | MASKED | no PMD correction | 14.27 | BAM | Shotgun (unfiltered) | 4,715,826 | 144,190 |
| UstIshim | MASKED | no PMD correction | 22.96 | BAM | Shotgun (unfiltered) | 4,729,941 | 192,944 |
| GOR001 | MASKED | REFINING | 7.55 | FASTQ | Shotgun | 4,718,681 | 35,167 |
| Mota | MASKED | REFINING | 9.57 | FASTQ | Shotgun | 4,671,490 | 137,567 |
| irk034 | MASKED | REFINING | 14.60 | FASTQ | Shotgun | 4,762,615 | 422,046 |
| mfo001 | MASKED | REFINING | 6.93 | FASTQ | Shotgun | 4,701,116 | 138,436 |
| saqqaq | MASKED | REFINING | 13.09 | FASTQ | Shotgun | 4,156,878 | 96,917 |
| Bon002 | MASKED | REFINING | 6.69 | FASTQ | Whole Genome Capture | 4,383,870 | 29,562 |
| prs013 | MASKED | REFINING | 4.68 | FASTQ | Whole Genome Capture | 4,512,973 | 4,658 |
| I0018 | MASKED | REFINING | 10.45 | BAM | 1240K Capture | 1,023,386 | 10,366 |
| I0585 | MASKED | REFINING | 19.11 | BAM | 1240K Capture | 2,102,549 | 19,394 |
| I0709 | MASKED | REFINING | 10.65 | BAM | 1240K Capture | 1,305,663 | 17,337 |
| I2452 | MASKED | REFINING | 8.74 | BAM | 1240K Capture | 836,319 | 13,042 |

| | | | | | | | |
|---|---|---|---|---|---|---|---|
| I4106 | MASKED | REFINING | 5.70 | BAM | 1240K Capture | 821,796 | 2,934 |
| Anzick.1 | MASKED | REFINING | 11.91 | BAM | Shotgun (filtered) | 4,588,077 | 167,847 |
| BOT2016 | MASKED | REFINING | 12.08 | BAM | Shotgun (filtered) | 4,750,124 | 197,868 |
| Yamnaya | MASKED | REFINING | 20.29 | BAM | Shotgun (filtered) | 4,752,494 | 419,303 |
| LBK | MASKED | REFINING | 14.27 | BAM | Shotgun (unfiltered) | 4,714,542 | 411,966 |
| UstIshim | MASKED | REFINING | 22.96 | BAM | Shotgun (unfiltered) | 4,728,992 | 543,495 |
| GOR001 | MASKED | TRIMMING | 7.55 | FASTQ | Shotgun | 4,702,184 | 35,167 |
| Mota | MASKED | TRIMMING | 9.57 | FASTQ | Shotgun | 4,652,646 | 137,567 |
| irk034 | MASKED | TRIMMING | 14.60 | FASTQ | Shotgun | 4,762,235 | 422,046 |
| mfo001 | MASKED | TRIMMING | 6.93 | FASTQ | Shotgun | 4,697,972 | 138,436 |
| saqqaq | MASKED | TRIMMING | 13.09 | FASTQ | Shotgun | 4,130,856 | 96,917 |
| Bon002 | MASKED | TRIMMING | 6.69 | FASTQ | Whole Genome Capture | 4,257,296 | 29,562 |
| prs013 | MASKED | TRIMMING | 4.68 | FASTQ | Whole Genome Capture | 4,395,349 | 4,658 |
| I0018 | MASKED | TRIMMING | 10.45 | BAM | 1240K Capture | 1,000,061 | 10,366 |
| I0585 | MASKED | TRIMMING | 19.11 | BAM | 1240K Capture | 2,060,493 | 19,394 |
| I0709 | MASKED | TRIMMING | 10.65 | BAM | 1240K Capture | 1,281,103 | 17,337 |
| I2452 | MASKED | TRIMMING | 8.74 | BAM | 1240K Capture | 820,712 | 13,042 |
| I4106 | MASKED | TRIMMING | 5.70 | BAM | 1240K Capture | 806,329 | 2,934 |
| Anzick.1 | MASKED | TRIMMING | 11.91 | BAM | Shotgun (filtered) | 4,532,077 | 167,847 |
| BOT2016 | MASKED | TRIMMING | 12.08 | BAM | Shotgun (filtered) | 4,745,429 | 197,868 |
| Yamnaya | MASKED | TRIMMING | 20.29 | BAM | Shotgun (filtered) | 4,749,158 | 419,303 |
| LBK | MASKED | TRIMMING | 14.27 | BAM | Shotgun (unfiltered) | 4,713,011 | 411,966 |

| | | | | | | | |
|---|---|---|---|---|---|---|---|
| UstIshim | MASKED | TRIMMING | 22.96 | BAM | Shotgun (unfiltered) | 4,728,021 | 543,495 |
| GOR001 | GRAPH | no PMD correction | 7.55 | FASTQ | Shotgun | 4,762,097 | 12,241 |
| Mota | GRAPH | no PMD correction | 9.57 | FASTQ | Shotgun | 4,739,777 | 48,387 |
| irk034 | GRAPH | no PMD correction | 14.60 | FASTQ | Shotgun | 4,770,122 | 150,413 |
| mfo001 | GRAPH | no PMD correction | 6.93 | FASTQ | Shotgun | 4,745,609 | 48,480 |
| saqqaq | GRAPH | no PMD correction | 13.09 | FASTQ | Shotgun | 4,516,002 | 31,042 |
| Bon002 | GRAPH | no PMD correction | 6.69 | FASTQ | Whole Genome Capture | 4,578,573 | 10,405 |
| prs013 | GRAPH | no PMD correction | 4.68 | FASTQ | Whole Genome Capture | 4,666,611 | 1,605 |
| I0018 | GRAPH | no PMD correction | 10.45 | BAM | 1240K Capture | 1,083,157 | 3,643 |
| I0585 | GRAPH | no PMD correction | 19.11 | BAM | 1240K Capture | 2,213,067 | 7,018 |
| I0709 | GRAPH | no PMD correction | 10.65 | BAM | 1240K Capture | 1,384,885 | 6,030 |
| I2452 | GRAPH | no PMD correction | 8.74 | BAM | 1240K Capture | 886,939 | 4,425 |
| I4106 | GRAPH | no PMD correction | 5.70 | BAM | 1240K Capture | 876,335 | 991 |
| Anzick.1 | GRAPH | no PMD correction | 11.91 | BAM | Shotgun (filtered) | 4,647,645 | 55,641 |
| BOT2016 | GRAPH | no PMD correction | 12.08 | BAM | Shotgun (filtered) | 4,762,686 | 70,033 |
| Yamnaya | GRAPH | no PMD correction | 20.29 | BAM | Shotgun (filtered) | 4,764,273 | 149,249 |
| LBK | GRAPH | no PMD correction | 14.27 | BAM | Shotgun (unfiltered) | 4,747,460 | 144,194 |
| UstIshim | GRAPH | no PMD correction | 22.96 | BAM | Shotgun (unfiltered) | 4,757,783 | 192,941 |
| GOR001 | GRAPH | REFINING | 7.55 | FASTQ | Shotgun | 4,757,722 | 35,167 |
| Mota | GRAPH | REFINING | 9.57 | FASTQ | Shotgun | 4,731,989 | 137,567 |
| irk034 | GRAPH | REFINING | 14.60 | FASTQ | Shotgun | 4,770,066 | 422,046 |

| | | | | | | | |
|---|---|---|---|---|---|---|---|
| mfo001 | GRAPH | REFINING | 6.93 | FASTQ | Shotgun | 4,743,824 | 138,436 |
| saqqaq | GRAPH | REFINING | 13.09 | FASTQ | Shotgun | 4,503,515 | 96,917 |
| Bon002 | GRAPH | REFINING | 6.69 | FASTQ | Whole Genome Capture | 4,506,443 | 29,562 |
| prs013 | GRAPH | REFINING | 4.68 | FASTQ | Whole Genome Capture | 4,610,554 | 4,658 |
| I0018 | GRAPH | REFINING | 10.45 | BAM | 1240K Capture | 1,061,341 | 10,366 |
| I0585 | GRAPH | REFINING | 19.11 | BAM | 1240K Capture | 2,168,413 | 19,394 |
| I0709 | GRAPH | REFINING | 10.65 | BAM | 1240K Capture | 1,359,111 | 17,337 |
| I2452 | GRAPH | REFINING | 8.74 | BAM | 1240K Capture | 870,354 | 13,042 |
| I4106 | GRAPH | REFINING | 5.70 | BAM | 1240K Capture | 858,784 | 2,934 |
| Anzick.1 | GRAPH | REFINING | 11.91 | BAM | Shotgun (filtered) | 4,610,604 | 167,847 |
| BOT2016 | GRAPH | REFINING | 12.08 | BAM | Shotgun (filtered) | 4,760,458 | 197,868 |
| Yamnaya | GRAPH | REFINING | 20.29 | BAM | Shotgun (filtered) | 4,762,571 | 419,303 |
| LBK | GRAPH | REFINING | 14.27 | BAM | Shotgun (unfiltered) | 4,746,696 | 411,966 |
| UstIshim | GRAPH | REFINING | 22.96 | BAM | Shotgun (unfiltered) | 4,757,311 | 543,495 |
| GOR001 | GRAPH | TRIMMING | 7.55 | FASTQ | Shotgun | 4,750,835 | 35,167 |
| Mota | GRAPH | TRIMMING | 9.57 | FASTQ | Shotgun | 4,721,957 | 137,567 |
| irk034 | GRAPH | TRIMMING | 14.60 | FASTQ | Shotgun | 4,769,993 | 422,046 |
| mfo001 | GRAPH | TRIMMING | 6.93 | FASTQ | Shotgun | 4,742,050 | 138,436 |
| saqqaq | GRAPH | TRIMMING | 13.09 | FASTQ | Shotgun | 4,488,627 | 96,917 |
| Bon002 | GRAPH | TRIMMING | 6.69 | FASTQ | Whole Genome Capture | 4,414,768 | 29,562 |
| prs013 | GRAPH | TRIMMING | 4.68 | FASTQ | Whole Genome Capture | 4,516,603 | 4,658 |
| I0018 | GRAPH | TRIMMING | 10.45 | BAM | 1240K Capture | 1,037,197 | 10,366 |
| I0585 | GRAPH | TRIMMING | 19.11 | BAM | 1240K Capture | 2,125,673 | 19,394 |

| | | | | | | | |
|---|---|---|---|---|---|---|---|
| I0709 | GRAPH | TRIMMING | 10.65 | BAM | 1240K Capture | 1,333,770 | 17,337 |
| I2452 | GRAPH | TRIMMING | 8.74 | BAM | 1240K Capture | 853,856 | 13,042 |
| I4106 | GRAPH | TRIMMING | 5.70 | BAM | 1240K Capture | 842,411 | 2,934 |
| Anzick.1 | GRAPH | TRIMMING | 11.91 | BAM | Shotgun (filtered) | 4,556,854 | 167,847 |
| BOT2016 | GRAPH | TRIMMING | 12.08 | BAM | Shotgun (filtered) | 4,757,754 | 197,868 |
| Yamnaya | GRAPH | TRIMMING | 20.29 | BAM | Shotgun (filtered) | 4,760,677 | 419,303 |
| LBK | GRAPH | TRIMMING | 14.27 | BAM | Shotgun (unfiltered) | 4,745,691 | 411,966 |
| UstIshim | GRAPH | TRIMMING | 22.96 | BAM | Shotgun (unfiltered) | 4,756,761 | 543,495 |

**Table S7:** Number of reads that mapped to linear, masked or graph genomes without mapping quality filtering (“No MAPQ filtering”) and with >30 mapping quality (“MAPQ > 30”). The sample ID’s in bold letters are the genomes available as FASTQ files.

| | **LINEAR** | | **MASKED** | | **GRAPH** | |
|---|---|---|---|---|---|---|
| **Sample ID** | **No MAPQ filtering** | **MAPQ > 30** | **No MAPQ filtering** | **MAPQ > 30** | **No MAPQ filtering** | **MAPQ > 30** |
| Anzick.1 | 498,389,271 | 496,271,671 | 498,056,319 | 494,062,850 | 486,279,290 | 477,696,119 |
| **Bon002** | 540,796,743 | 262,288,413 | 541,279,502 | 260,158,198 | 475,435,102 | 317,112,605 |
| BOT2016 | 455,858,162 | 455,174,186 | 455,115,983 | 453,426,390 | 452,413,987 | 447,810,130 |
| **GOR001** | 390,278,032 | 302,841,210 | 389,621,136 | 301,452,493 | 374,017,341 | 315,122,438 |
| I0018 | 23,117,836 | 19,165,269 | 23,116,698 | 19,126,565 | 22,074,066 | 19,112,729 |
| I0585 | 61,084,061 | 50,549,135 | 61,077,204 | 50,421,948 | 57,983,490 | 50,118,033 |
| I0709 | 30,365,941 | 23,996,671 | 30,359,353 | 23,927,362 | 28,564,603 | 23,902,478 |
| I2452 | 17,924,867 | 14,430,181 | 17,922,144 | 14,398,686 | 16,894,126 | 14,394,400 |
| I4106 | 15,085,187 | 11,253,341 | 15,082,847 | 11,220,225 | 14,051,602 | 11,232,060 |
| **irk034** | 495,966,990 | 432,374,861 | 495,048,585 | 430,918,725 | 512,301,500 | 456,308,204 |
| LBK | 703,869,504 | 627,129,925 | 703,498,280 | 625,933,531 | 677,038,587 | 625,491,111 |
| **mfo001** | 388,863,436 | 301,366,311 | 388,885,674 | 300,790,666 | 370,435,869 | 314,126,325 |
| **Mota** | 522,418,413 | 445,246,848 | 522,125,302 | 444,173,147 | 566,005,792 | 475,974,923 |
| **prs013** | 327,195,151 | 248,292,569 | 327,138,442 | 247,542,464 | 318,491,246 | 261,880,277 |
| **saqqaq** | 953,857,718 | 689,180,685 | 950,874,032 | 685,869,000 | 947,801,336 | 788,713,640 |
| UstIshim | 1,339,153,486 | 1,136,973,730 | 1,338,791,590 | 1,135,063,885 | 1,252,364,327 | 1,132,958,000 |
| Yamnaya | 881,666,147 | 880,133,525 | 880,997,137 | 877,385,333 | 873,296,099 | 863,898,144 |

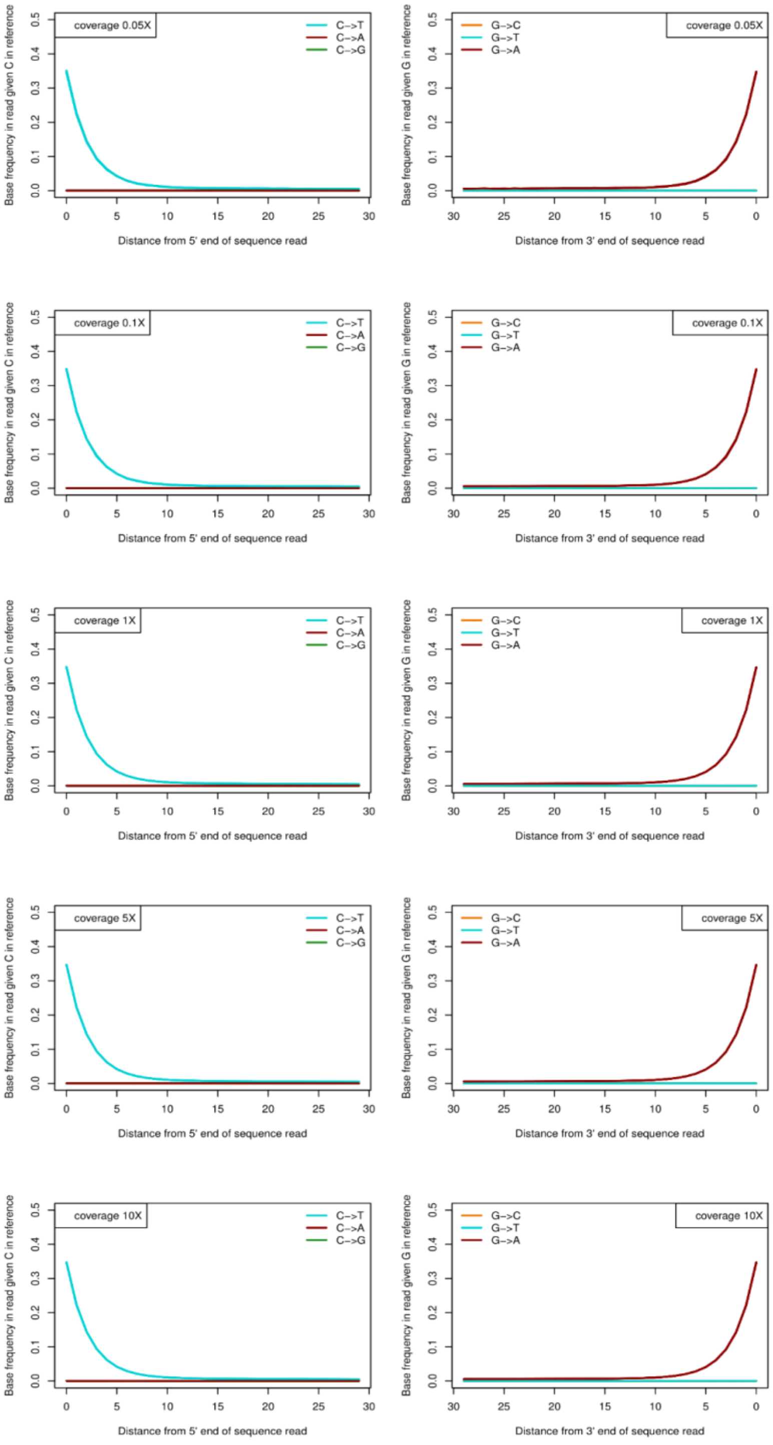


**Figure S1:** Post-mortem damage patterns in ancient DNA-like simulated data after mapping to the linear reference genome. The y-axis shows different mutation rates and the x-axis shows the position from the read 5’-end (left panels) or the 3’-end (right panels).

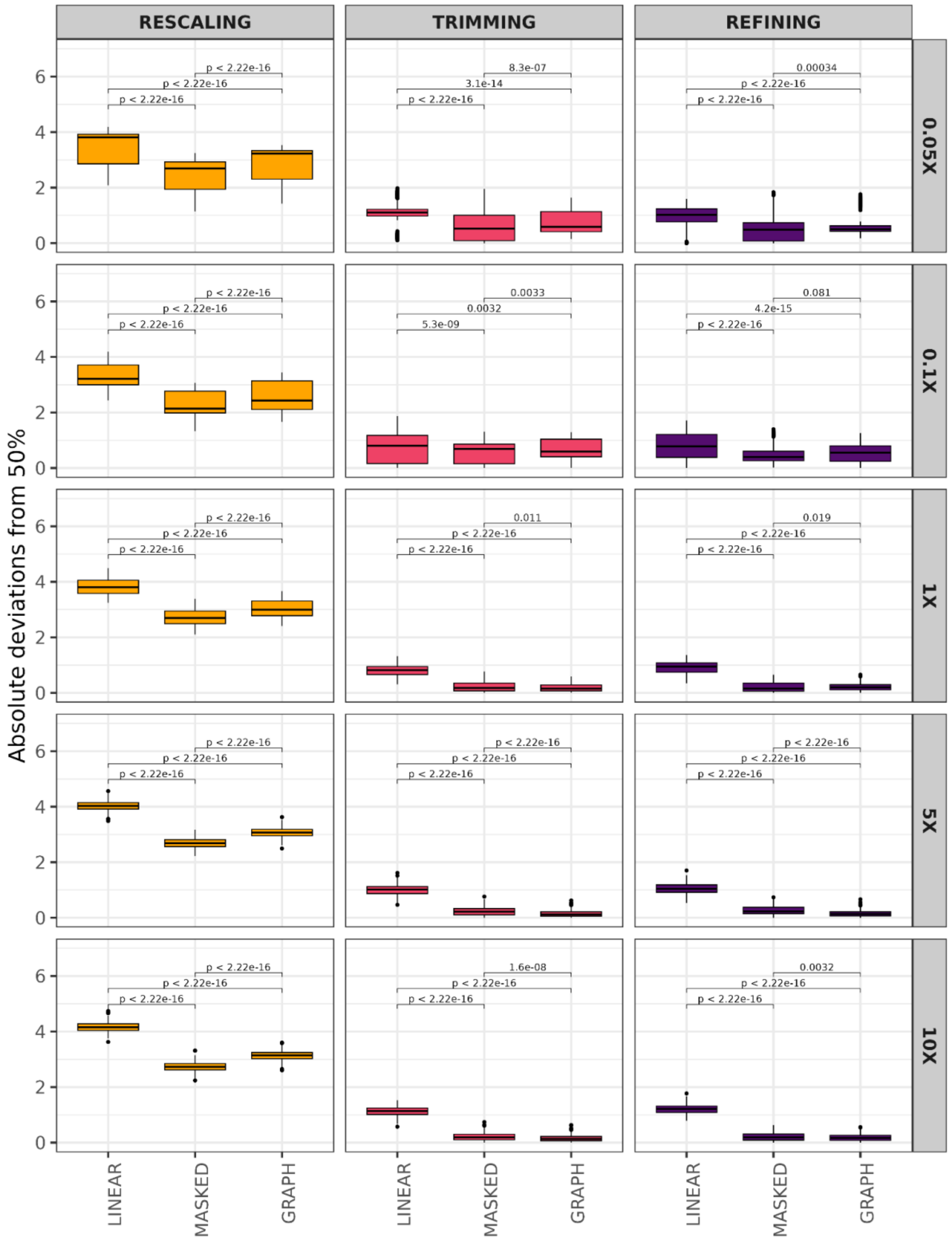


**Figure S2:** Absolute deviations from 50% on heterozygote sites on aDNA-like simulated data using the RESCALING, TRIMMING and REFINING strategies for PMD correction between different mapping strategies. The distributions were obtained using 100 replicates at different coverages (rows). The p-values were calculated on pairwise comparisons using the Mann-Whitney U test.

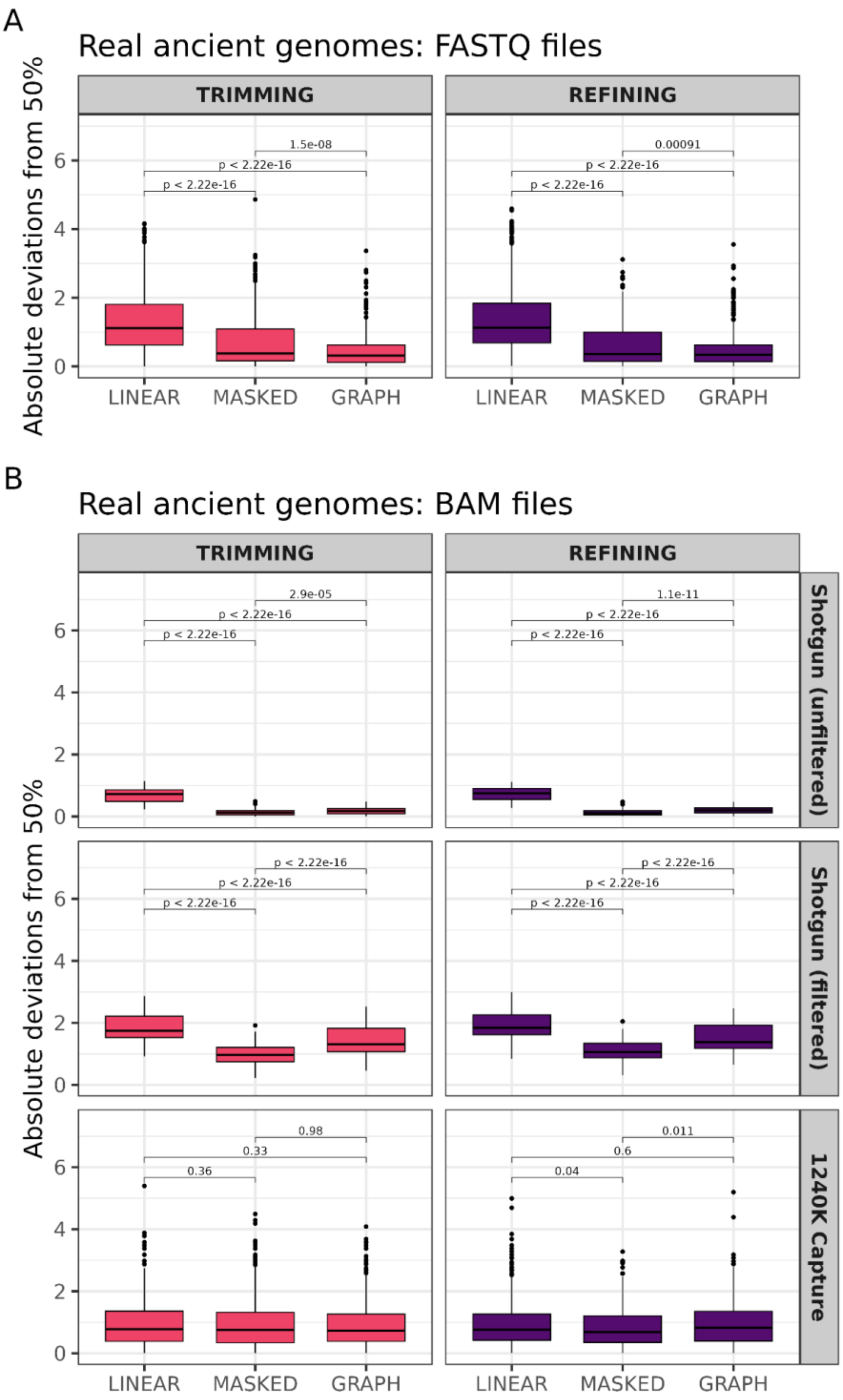


**Figure S3:** Absolute deviations from 50% on heterozygote sites on published ancient genomes using the “TRIMMING” and “REFINING” strategies for PMD correction between different mapping strategies. The distributions were obtained using 100 replicates for each sample. Panel A shows the results for ancient genomes with available raw FASTQ files, and Panel B shows results for ancient genomes published with already processed BAM files. The p-values were calculated on pairwise comparisons using the Mann-Whitney U test.

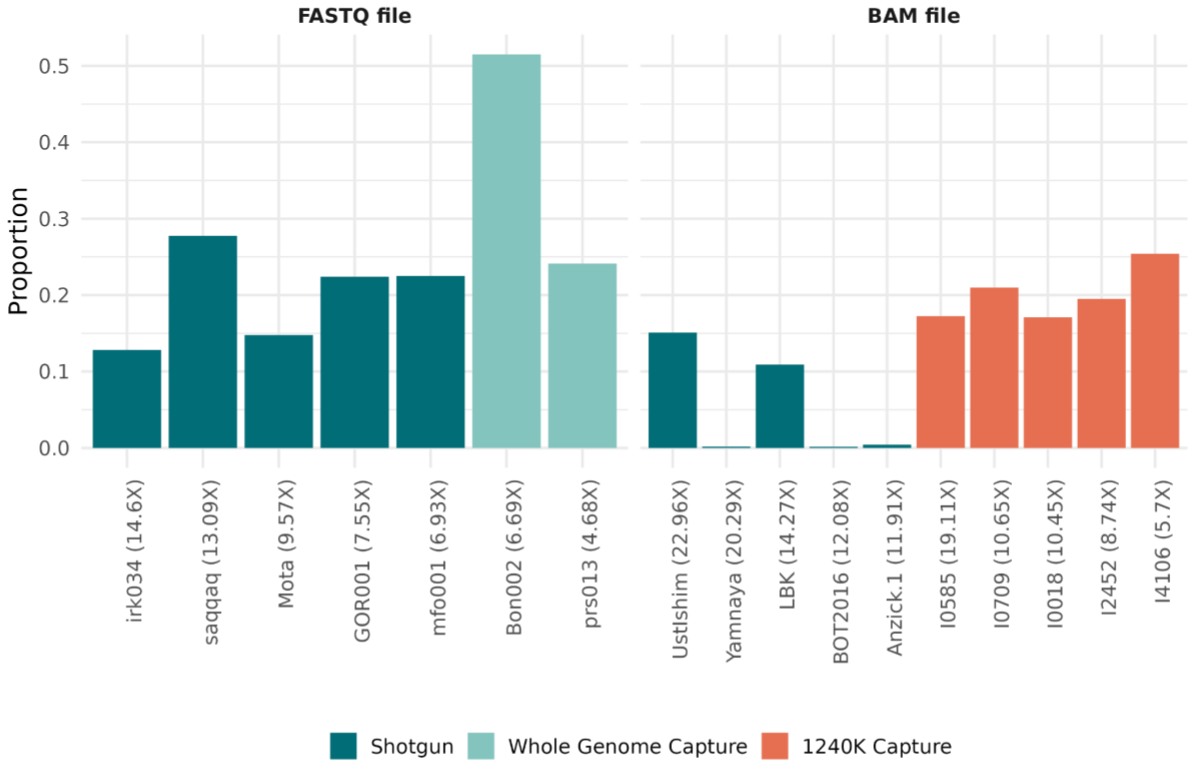


**Figure S4:** The proportion of the reads with mapping quality (MAPQ) <30 for each sample after aligning them to linear human reference genome. The published BAM files with practically no reads with MAPQ <30 can be assumed to be already quality-filtered.

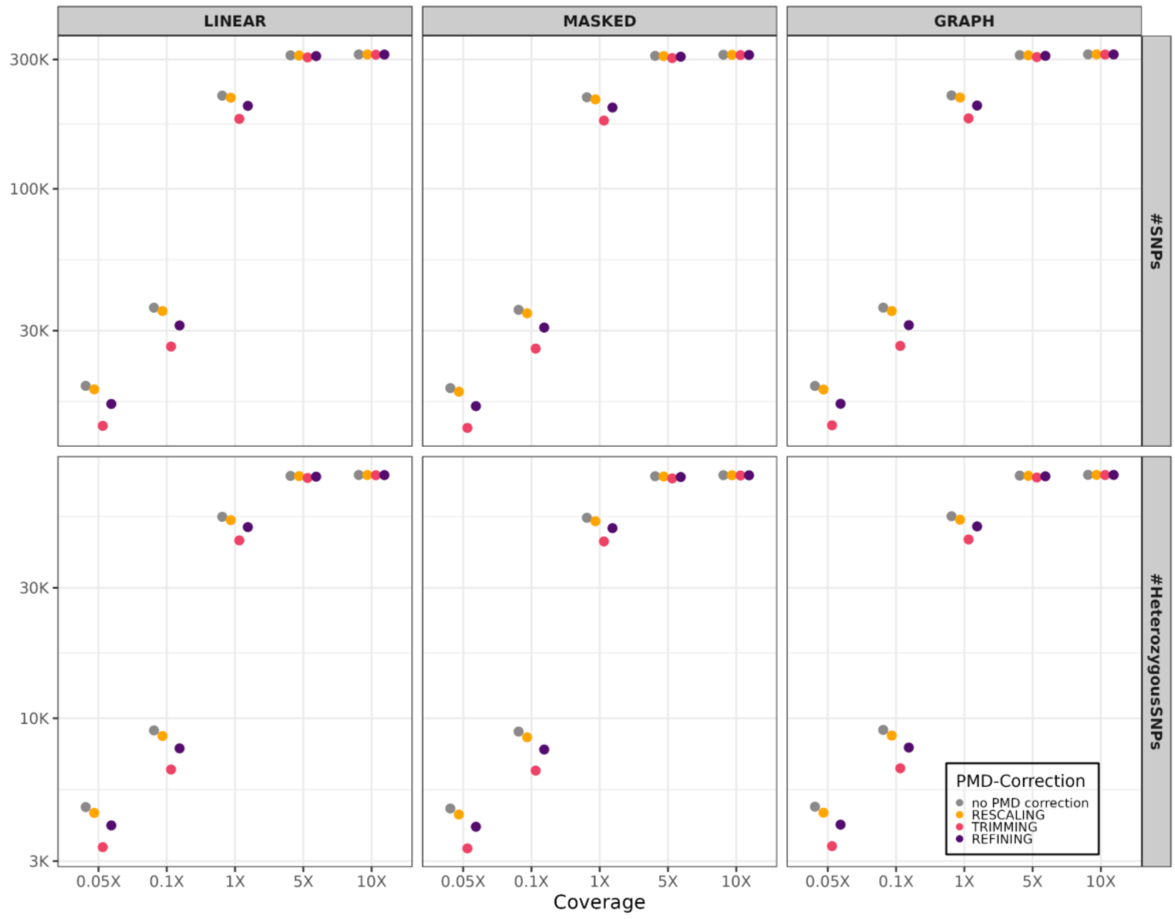


**Figure S5:** The average number of SNP genotyped for aDNA-like simulated genomes at different coverages and PMD correction strategies. The numbers were calculated after genotyping all targeted SNPs (n=313,747) in the upper panel or genotyping heterozygous positions (n=77,841) in the lower panel. Genotyping was performed with pileupCaller. There are 5 runs for each coverage, and the average is shown. Because the number of the genotyped SNPs was highly similar, standard errors are not visible.

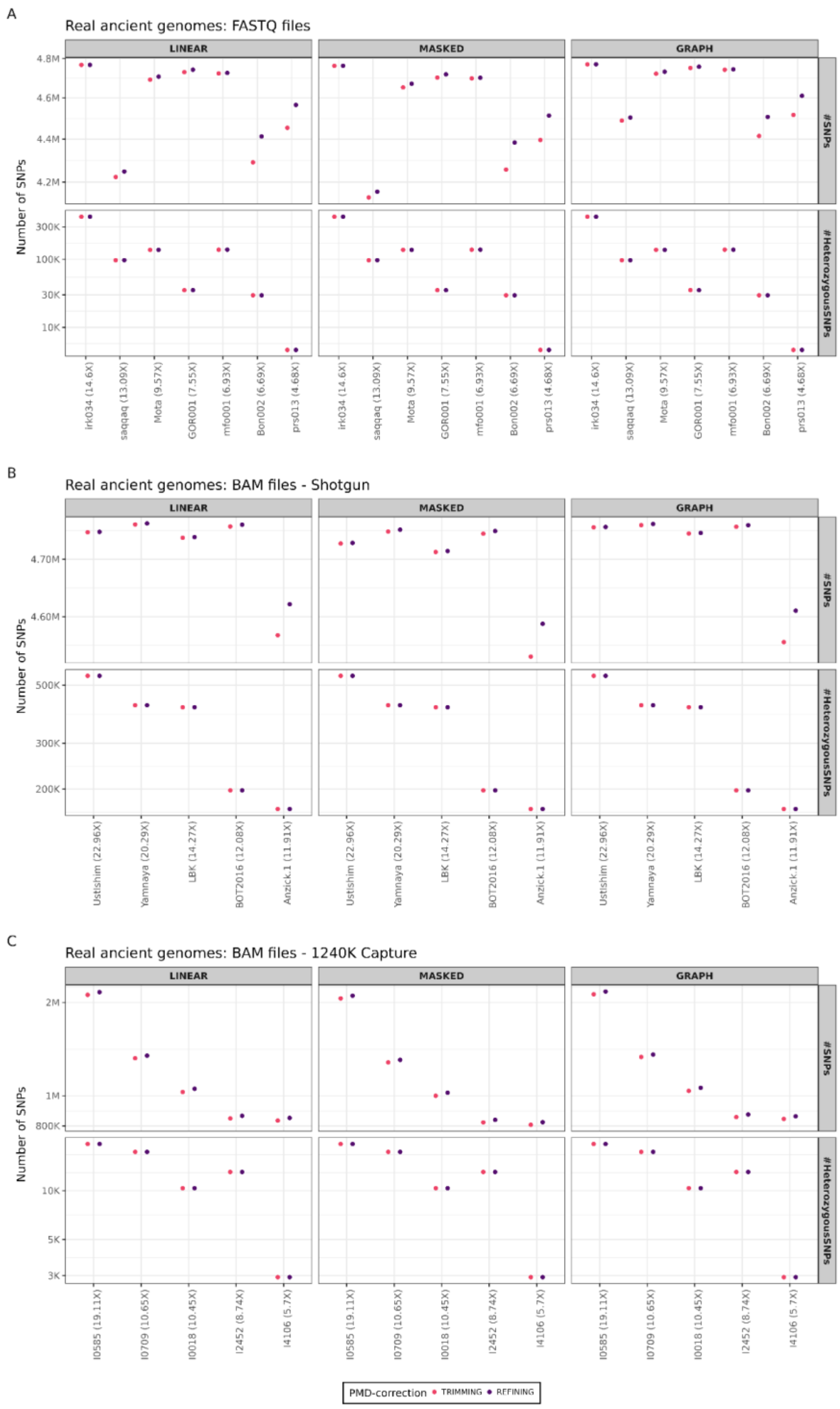


**Figure S6:** The average number of SNP genotyped in real ancient genomes and using “TRIMMING” or “REFINING” as PMD correction strategies. The numbers were calculated after genotyping all targeted SNPs in the upper panels or genotyping heterozygous positions in the lower panels. Heterozygous positions were defined as positions where alternative allele frequencies were 25-75% and had a minimum of 10 reads (Table S4). Genotyping was performed with pileupCaller. Panels A, B and C show shotgun genomes available as FASTQ files, shotgun genomes available as BAM files, and 1240K capture genomes available as BAM files, respectively. The genome sample IDs are shown above panels.

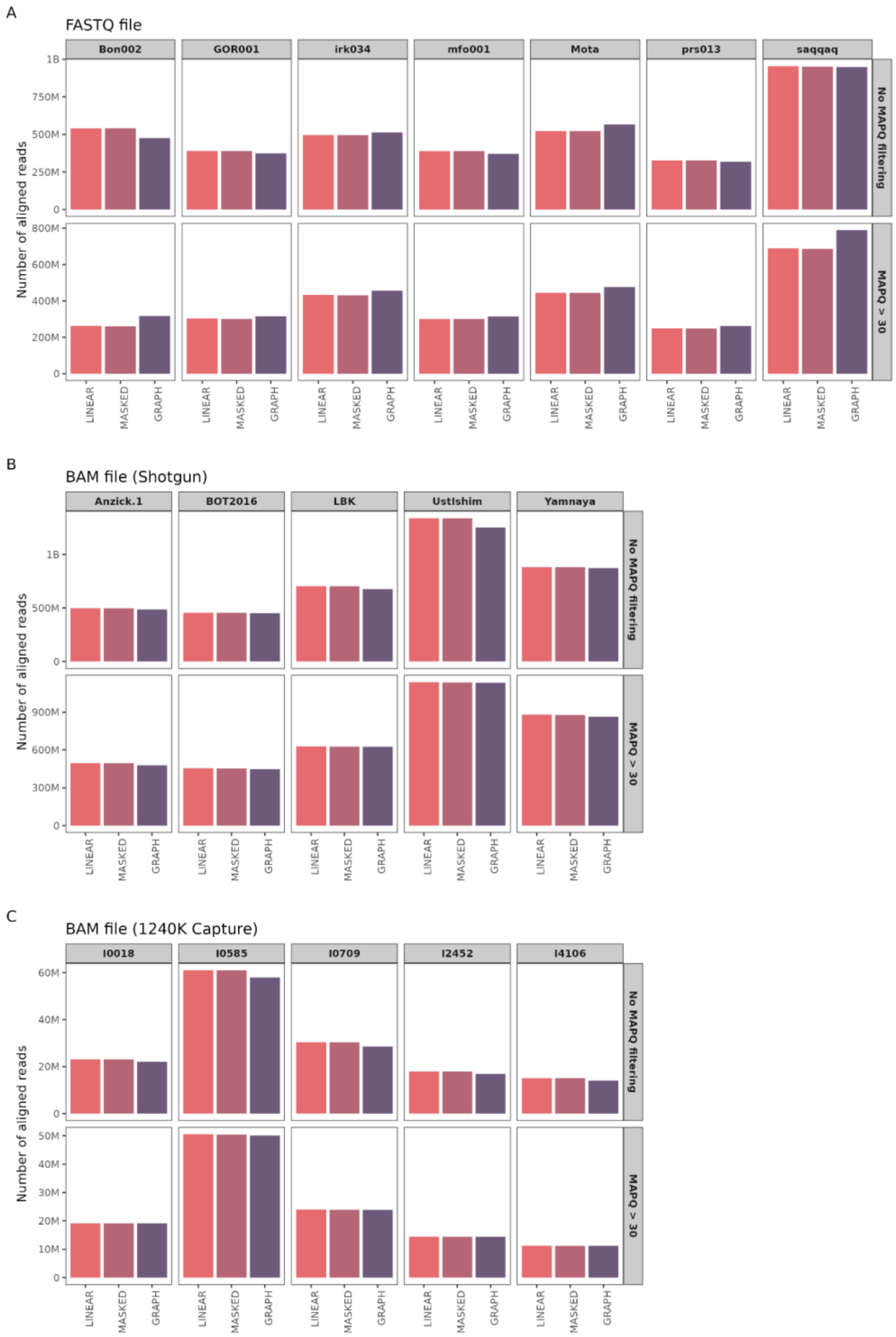


**Figure S7:** Comparing the number of aligned reads under three different mapping strategies, shown as x-axis labels. The figure shows the number of reads without (upper panels) and with mapping quality filters (lower panels). Panels A, B and C show shotgun genomes available as FASTQ files, shotgun genomes available as BAM files, and 1240K capture genomes available as BAM files, respectively. The genome sample IDs are shown above panels.

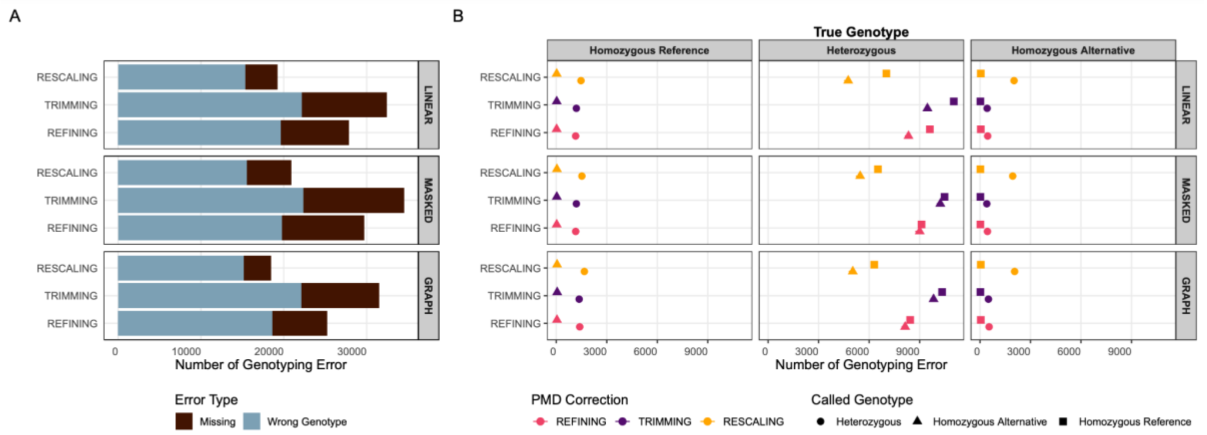


**Figure S8:** Performance of different alignment strategies and PMD correction strategies on simulated aDNA-like genomes with 5X coverage. (A) The proportion of genotyping errors and missingness. (B) The frequency of the type of genotyping errors for each PMD correction strategy.

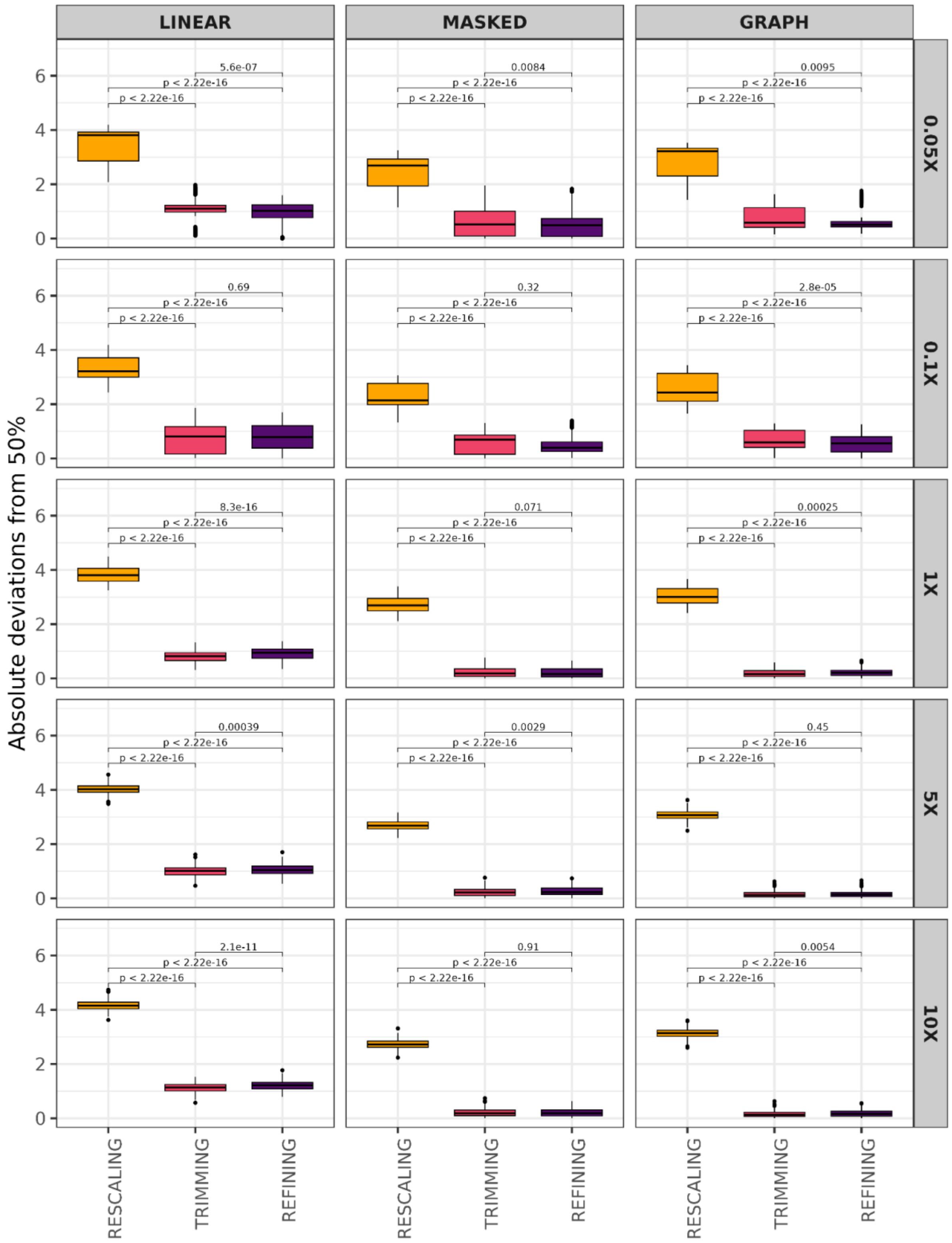


**Figure S9:** Absolute deviations from 50% on heterozygote sites on aDNA-like simulated data using the “RESCALING”, “TRIMMING” and “REFINING” strategies for PMD correction and using all three mapping strategies. The distributions were obtained using 100 replicates at different coverages (rows). The p-values were calculated on pairwise comparisons using the Mann-Whitney U test.

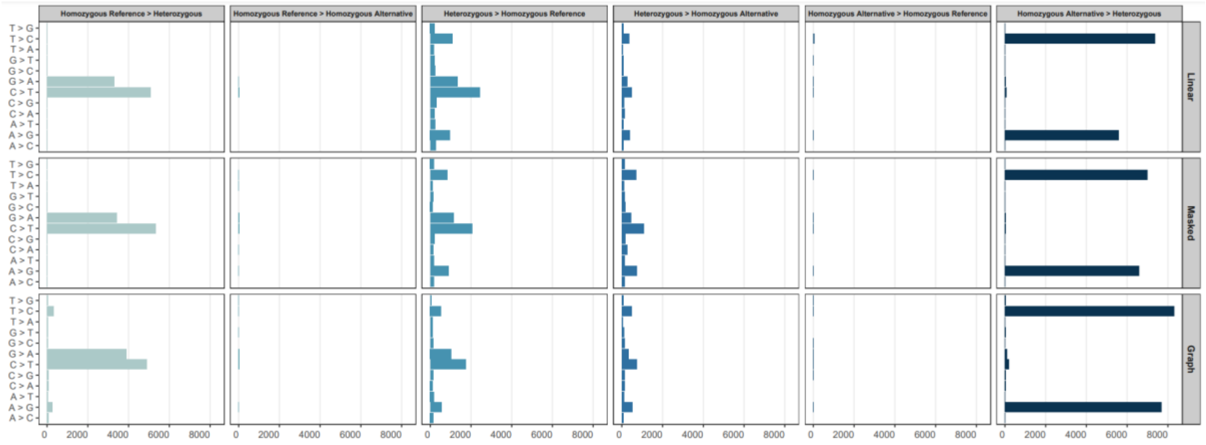


**Figure S10:** Distributions of different types of genotyping error introduced using the "RESCALING" strategy on simulated aDNA-like genomes with 10X coverage. The results were calculated using three alternative alignment strategies, shown on the right side. The panel titles show errors; e.g. "Homozygous Reference > Heterozygous" indicates a real homozygous reference genotype called heterozygous.

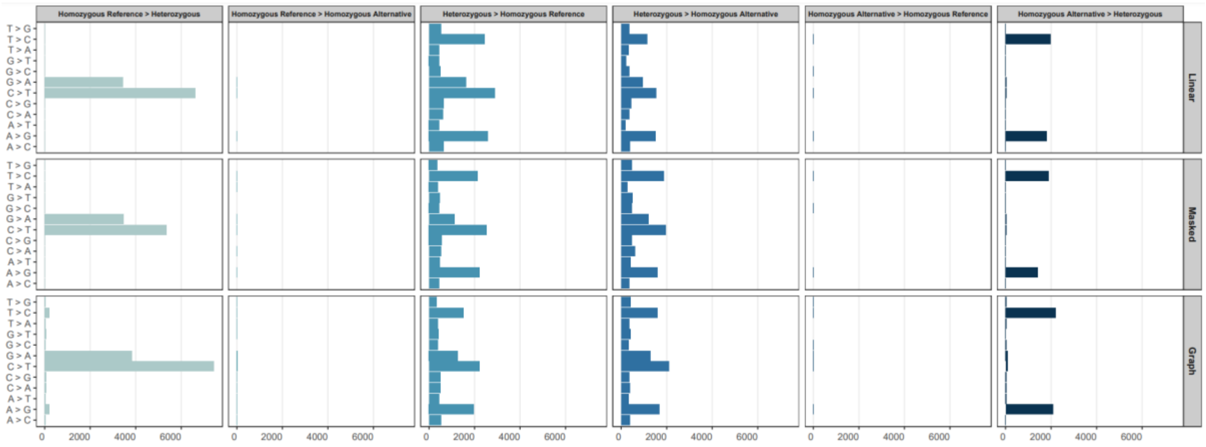


**Figure S11:** Distributions of different types of genotyping error introduced using the "TRIMMING" strategy on simulated aDNA-like genomes with 10X coverage. The results were calculated using three alternative alignment strategies, shown on the right side. The panel titles show errors; e.g. "Homozygous Reference > Heterozygous" indicates a real homozygous reference genotype called heterozygous.

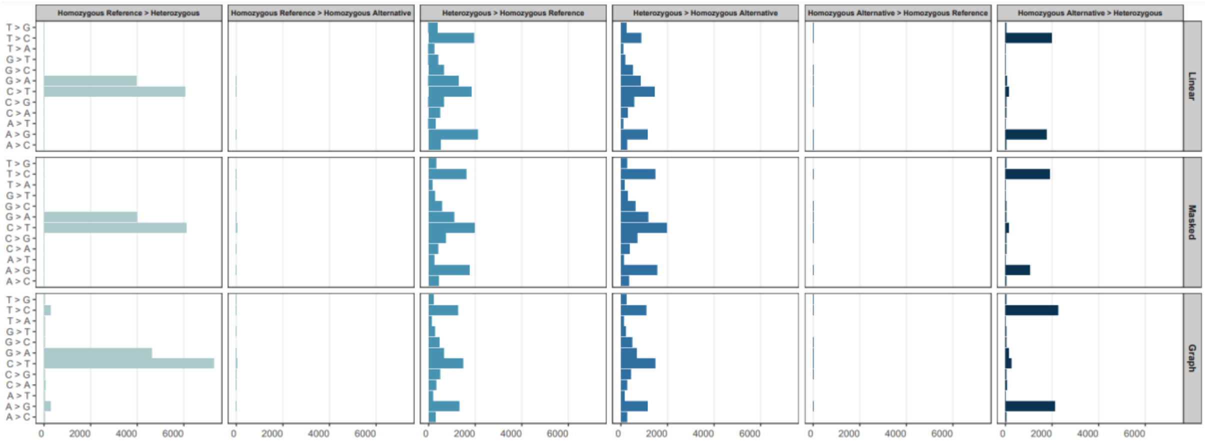


**Figure S12:** Distributions of different types of genotyping error introduced using the "REFINING" strategy on simulated aDNA-like genomes with 10X coverage. The results were calculated using three alternative alignment strategies, shown on the right side. The panel titles show errors; e.g. "Homozygous Reference > Heterozygous" indicates a real homozygous reference genotype called heterozygous.

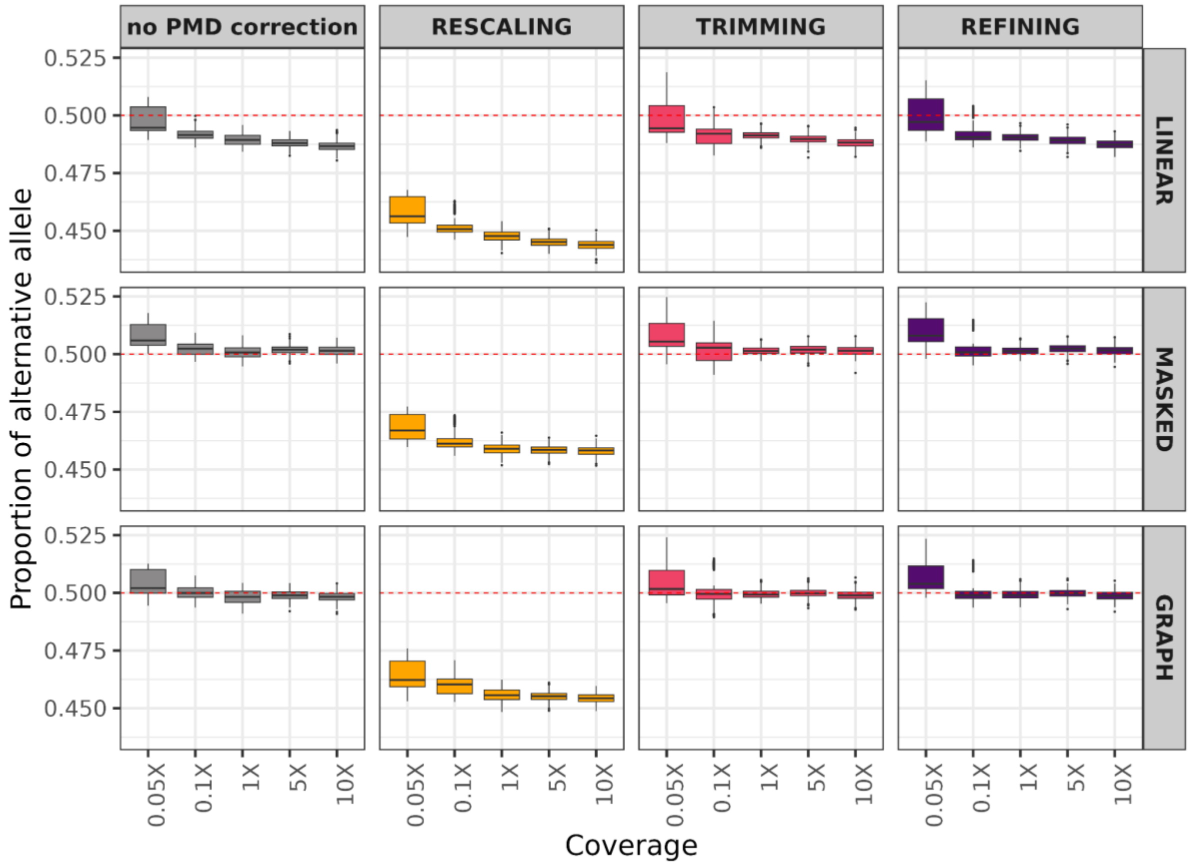


**Figure S13:** Comparing reference bias in simulated aDNA-like genomes at transition SNPs using different alignment strategies (rows, indicated on the right) and PMD correction strategies (columns, indicated in the panel title) and different coverages (indicated as x-axis labels). The plot shows the proportion of alternative alleles after randomly selecting one allele from heterozygote sites 100 times by pileupCaller. Here we only considered transition SNPs (see Figure 3). The reason for the relationship between reference bias and coverage in this data is not obvious to us.

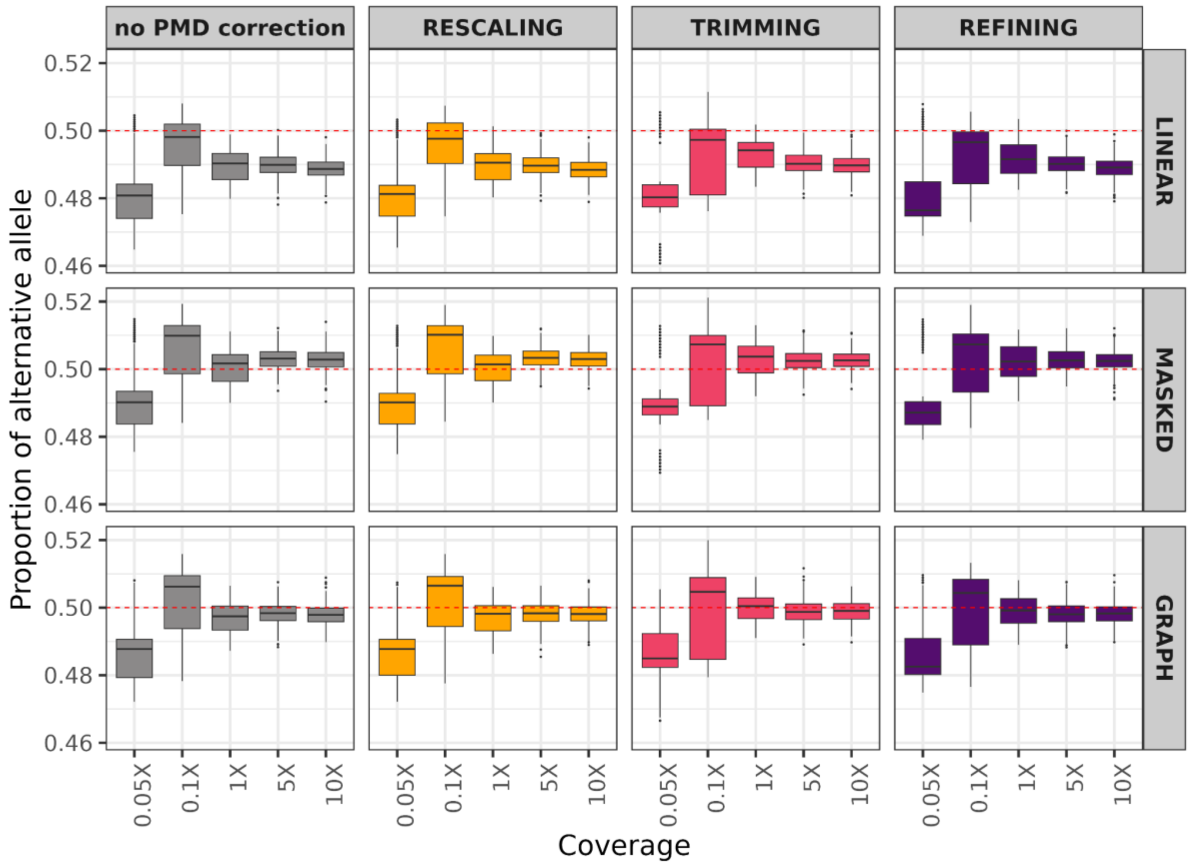


**Figure S14:** Comparing reference bias in simulated aDNA-like genomes at transversion SNPs using different alignment strategies (rows, indicated on the right) and PMD correction strategies (columns, indicated in the panel title) and different coverages (indicated as x-axis labels). The plot shows the proportion of alternative alleles after randomly selecting one allele from heterozygote sites 100 times by pileupCaller. Here we only considered transversion SNPs (see Figure 3). The reason for the higher reference bias at 0.05X is not obvious to us.

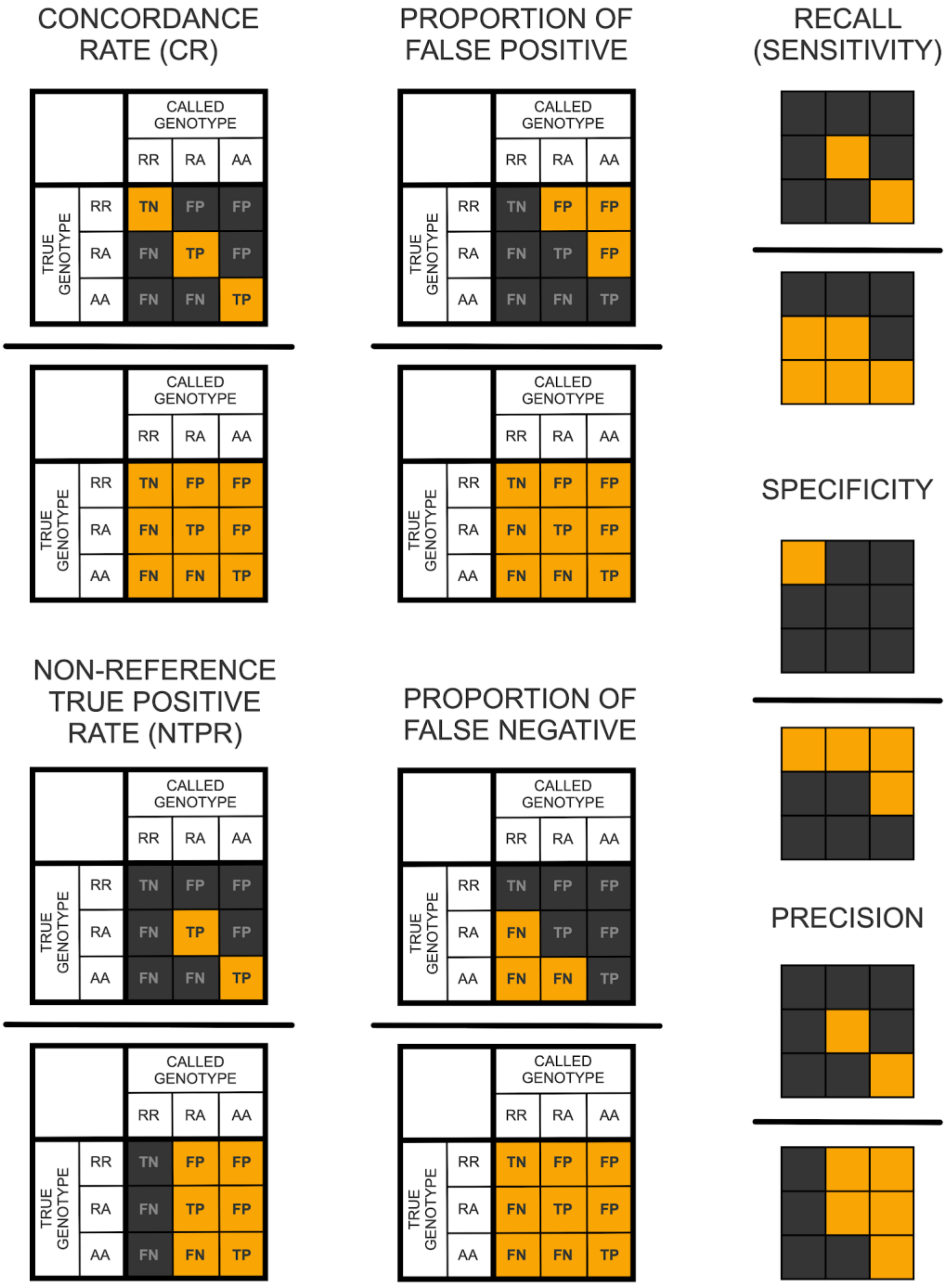


**Figure S15:** Genotype concordance measures used for Figure 4. All combinations of the genotyped positions [182,515 homozygous reference (RR), 53,391 homozygous alternative (AA), 77,841 heterozygous (RA)] were used for simulation. The panels show how the concordance rate (CR), false negative rate (FNR), false positive rate (FPR), non-reference true positive rate (NTPR), recall (or sensitivity), specificity and precision are calculated with the alternative allele as a pivot. The horizontal lines stand for division. The yellow-coloured cells are the frequencies included in the calculation for each statistic (e.g. CR is calculated by summing TN and the two TP frequencies and dividing by the total frequency of all observations). We also calculated the F-score, which is the harmonic mean of precision and recall values, by the following formula: 2 x [(Precision x Recall) / (Precision + Recall)]. TN: True Negative, TP: True Positive, FP: False Positive, FN: False Negative. The figure was adopted from Kishikawa et al., 2019.

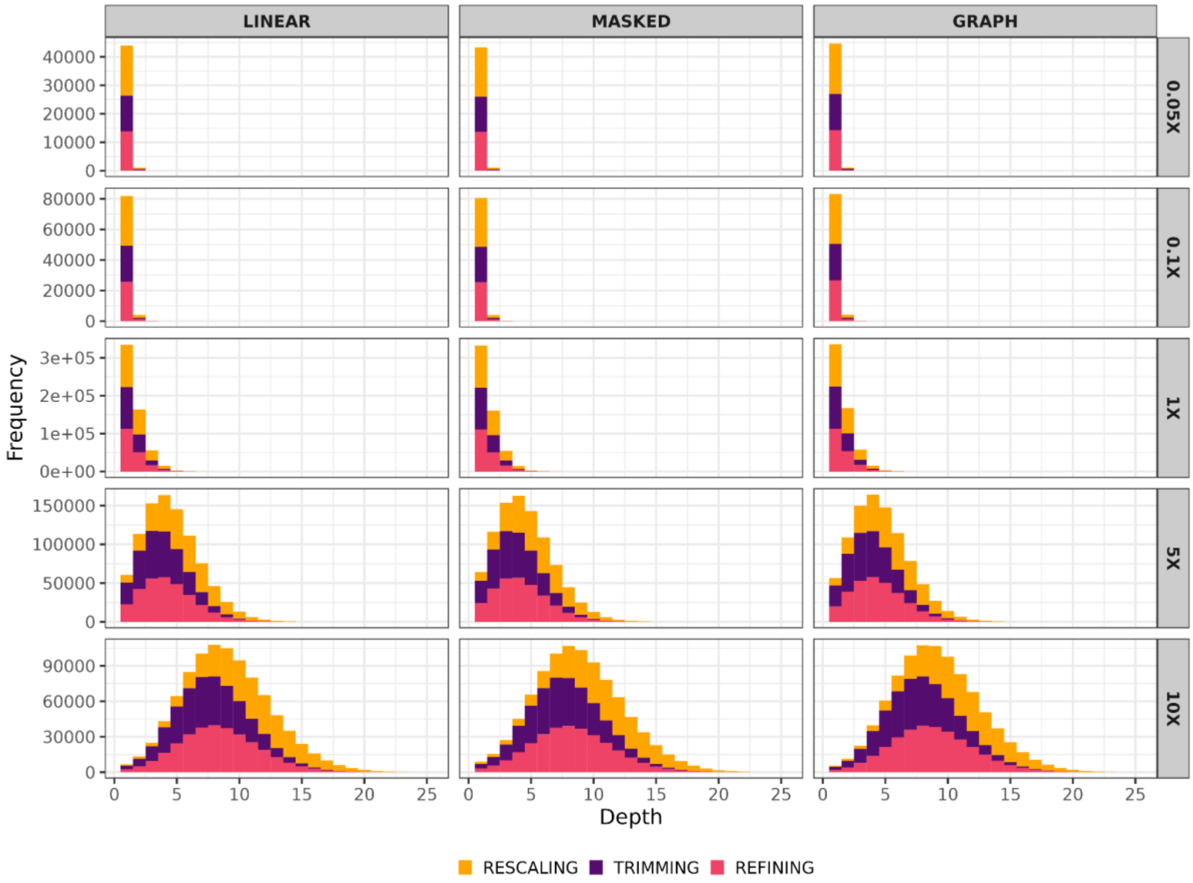


**Figure S16:** Depth-of-coverage at variable positions used for genotyping in aDNA-like simulated genomes at different coverages (rows), using three different alignment strategies (columns), and using three different PMD correction strategies (colours). The per variant coverage values were calculated based on '*samtools mpileup*' results, which were later used for creating pseudo-haploid genotypes (Methods).

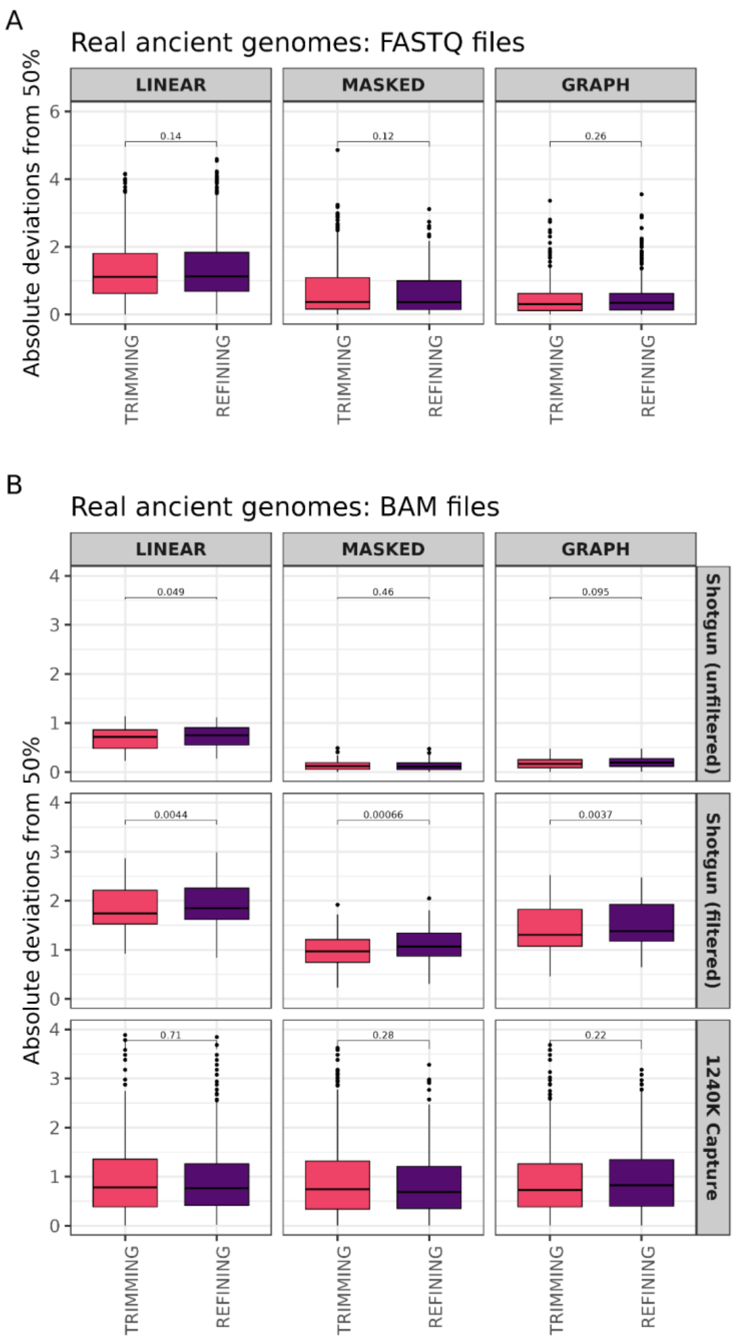


**Figure S17:** Absolute deviations from 50% on heterozygote sites on published ancient genomes using the "TRIMMING" and "REFINING" strategies for PMD correction and using all three mapping strategies. The distributions were obtained using 100 replicates for each sample. Panel A shows the results for ancient genomes with available raw FASTQ files, and Panel B shows results for ancient genomes published with already processed BAM files. The p-values were calculated on pairwise comparisons using the Mann-Whitney U test. The reason why "REFINING" performs slightly worse than "TRIMMING" in quality filtered shotgun data is not obvious to us.

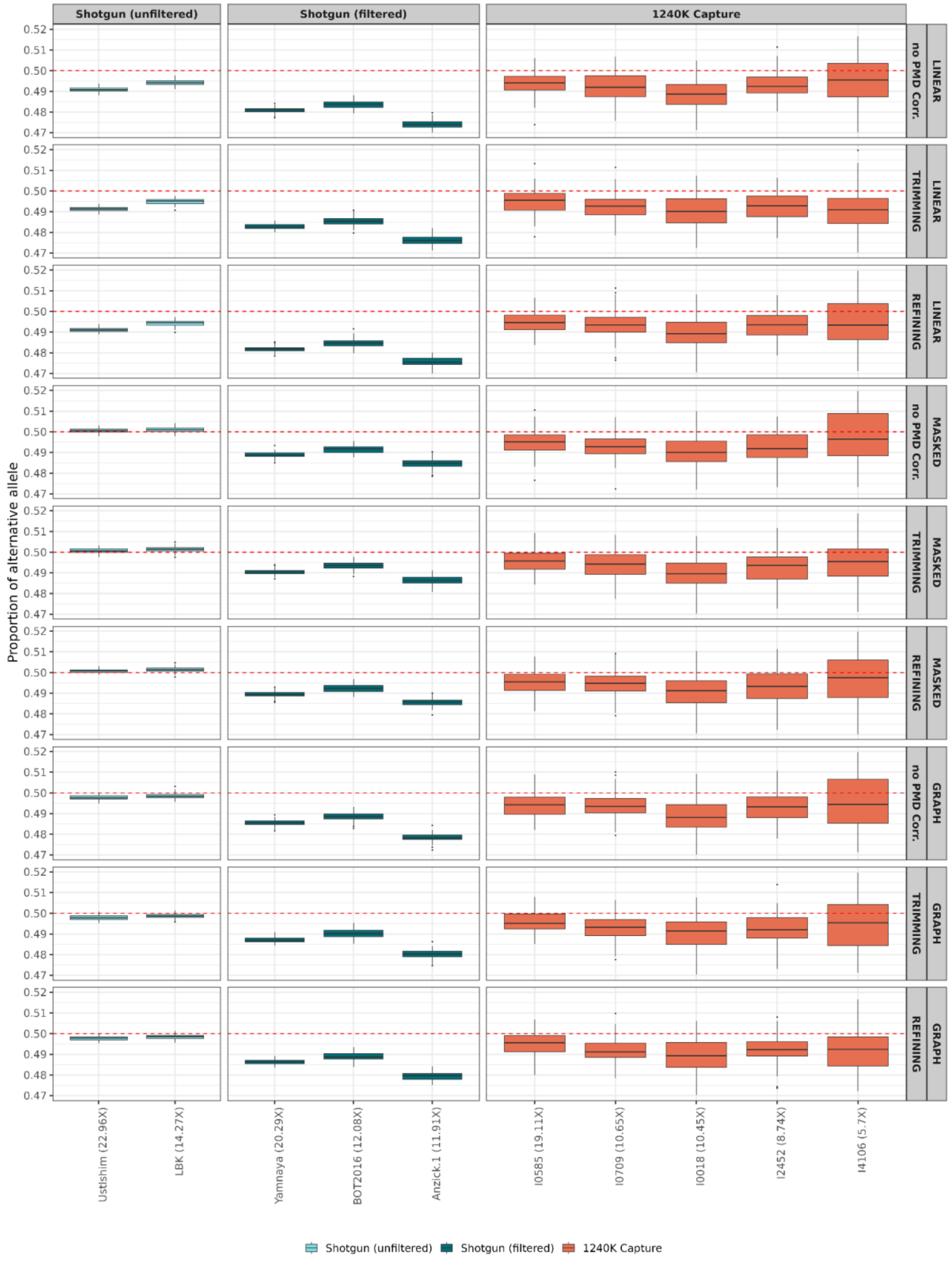


**Figure S18:** Comparing reference bias in published ancient genomes only published as BAM files using different alignment strategies and PMD correction strategies (in this case either “TRIMMING” or “REFINING”). The plot shows the proportion of alternative alleles after randomly selecting one allele from heterozygote sites 100 times by pileupCaller.

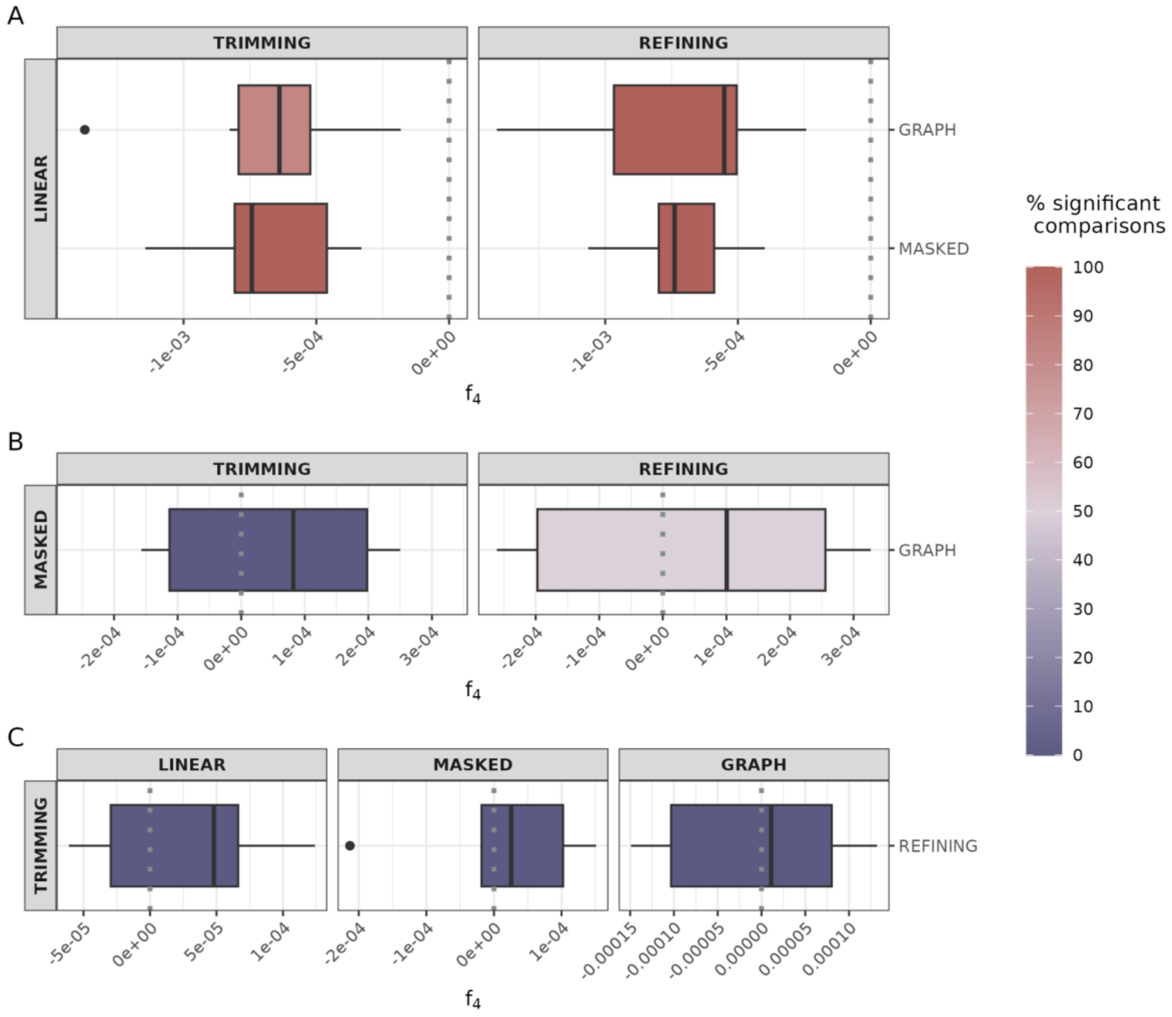


**Figure S19:** Reference bias measured using $f_4$ tests. Results from the model **A)** $f_4$(Chimp, Human Reference Genome; Ind1_LINEAR, Ind1_MASKED|GRAPH), **B)** $f_4$(Chimp, Human Reference Genome; Ind1_MASKED, Ind1_GRAPH) for both PMD correction strategies and **C)** $f_4$(Chimp, Human Reference Genome; Ind1_TRIMMED, Ind1_REFINED) for all mapping strategies by using ancient genomes that only BAM files available. The colour gradient from blue to red represents the fraction of comparisons that are nominally significant (|Z|>3).
